# Germline stem cell isolation, lineage tracing, and aging in a protochordate

**DOI:** 10.1101/2025.08.31.673173

**Authors:** Tom Levy, Chiara Anselmi, Katherine J. Ishizuka, Tal Gordon, Yotam Voskoboynik, Erin McGeever, Angela M. Detweiler, Liron Levin, Karla J. Palmeri, Daniel Dan Liu, Rahul Sinha, Benjamin F. Ohene-Gambill, Tal Raveh, Maurizio Morri, Virginia Vanni, Lucia Manni, Debashis Sahoo, Norma F. Neff, Benyamin Rosental, Irving L. Weissman, Ayelet Voskoboynik

## Abstract

Germline stem cells (GSCs), the source of gametes, are the only stem cells capable of passing genes to future generations making them cellular units of natural selection. Yet, factors regulating their fitness, and thus governing GSC competition, which exist in both protochordates and mammals, remain poorly understood. We study how aging affects GSC fitness in the protochordate *Botryllus schlosseri*, an evolutionary crosspoint between invertebrates and vertebrates. GSCs were isolated using flow cytometry and characterized by single-cell RNA sequencing, supported by a new PacBio genome assembly. GSC transplantation assays confirmed their function, further validated through a transgenerational lineage tracing strategy combining membrane-labeling dyes and single-cell transcriptomic analysis. Leveraging our method to isolate them, we uncover significant age-related changes in GSCs, while spermatids and sperm show minimal differences, suggesting that reproductive aging is influenced primarily by GSCs rather than by gametes. Reduced expression of markers like *DDX4* and *PIWIL1* in aged GSCs mirrors trends in mammalian datasets, suggesting conserved age-associated molecular changes in GSCs across chordate evolution. This study provides new techniques that lay the foundation to investigate further drivers of GSC fitness and highlights fertility-related genes as promising targets for therapies to preserve reproductive health.

## Introduction

Age-related decline in stem cell function and number within a given tissue can adversely affect tissue homeostasis^1^. This aging process may result from cell-intrinsic changes^2^, deterioration of niche support^1,3^ or both. While stem cells are responsible for the development and regeneration of tissues and organs, GSCs are the source of gametes and, unlike other stem cells, transmit their genomes to future generations via sexual reproduction. For this genetic transmission to occur, an organism must remain fertile. Therefore, it is highly important to investigate factors such as GSC aging, which may impair their fitness and potentially lead to reduced fertility^4–6^.

Research on the protochordate colonial tunicate *B. schlosseri* has significantly advanced our understanding of fundamental principles in stem cell biology, including stem cell competition at the germline level^7–11^ and stem cell aging^12^. GSC competition occurs when adjacent *Botryllus* colonies fuse blood vessels and create chimeras. In these chimeras, GSCs from one colony outcompete those from the other, resulting in gonads derived only from the ‘winner’ colony, which determines the genetic contribution to the next generation through sexual reproduction^7–11^. The germline winner outcome is heritable among independent individuals tested against loser colonies^9,10,13,14^. Since winner phenotypes are inherited across sexually reproductive generations, it is conceivable that winner phenotypes derive from genetic alleles. The phenomenon of GSC competition has been also demonstrated in mouse testes, but the controlling mechanisms, as in *Botryllus*, remain a mystery^15^. In addition to sexual reproduction, *Botryllus* colonies expand through weekly asexual budding^16^ (Fig. 1n). This unique process results in the coexistence of three asexual blastogenic generations within the same colony: adult zooids (first generation), primary buds (second generation), and secondary buds (third generation). During each weekly blastogenic cycle, adult zooids are resorbed and replaced by developing buds. Consequently, a new gonad forms weekly from GSCs in the secondary bud and matures in the adult zooid two weeks later, enabling continuous sexual reproduction. This blastogenic process is mediated by stem cells that migrate from resorbed zooids to the developing buds and create all the tissues and organs in subsequent asexual generations^11,16–18^. While the stem cell basis of this process was initially inferred from studies demonstrating stem cell competition in *Botryllus* natural chimeras^9,10,13,14^, it was further supported by prospective stem cell isolation, lineage tracing, and transplantation studies showing that circulating stem cells colonize developing buds and drive de novo organogenesis, whereas differentiated mature cells fail to engraft^11,18^. However, despite this transgenerational stem cell-mediated tissue renewal, aging colonies develop morphological, neural, and molecular defects over successive asexual generations^19–21^. Thus, because stem cells are suggested to persist throughout the colony’s lifespan, and age as they move across asexual generations, colony aging might reflect the aging of its stem cells^12^. Colony aging in *Botryllus* should therefore be distinguished from vertebrate organismal aging. Unlike vertebrate aging, which occurs in a single persistent individual, a *Botryllus* colony undergoes lifelong cyclical asexual replacement of its zooids while maintaining long-lived stem-cell lineages. This makes *B. schlosseri* a unique model for studying gonadal development at the cellular level and GSC aging.

**Fig. 1:**
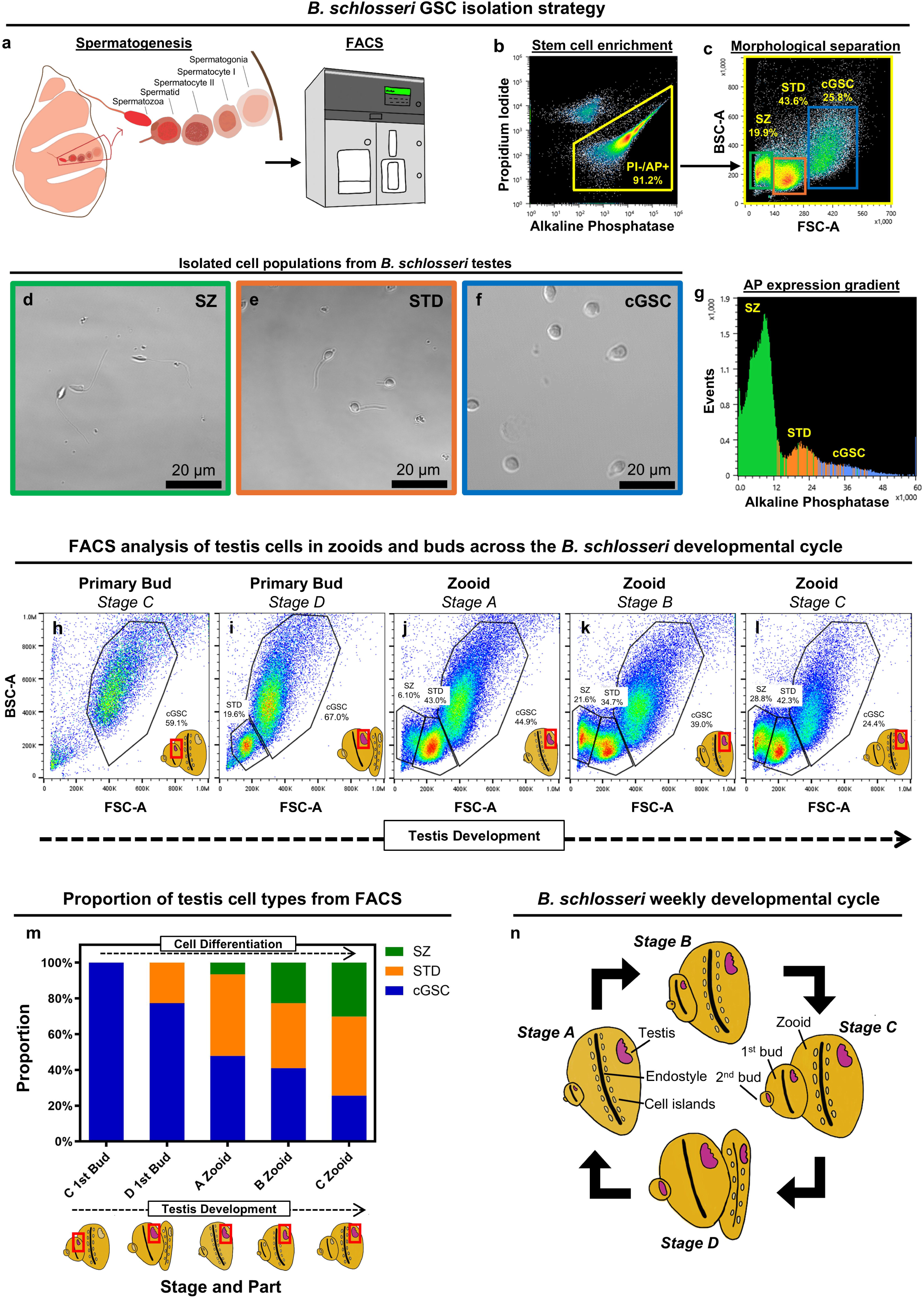
Identification of *B. schlosseri* spermatogenic germline cell types and cellular dynamics of testis development. **a,** Cell types identified during spermatogenesis. Spermatogonia are located at the lobe’s periphery, followed by primary and secondary spermatocytes and spermatids (STD). Fully mature sperm (spermatozoa; SZ) are found in the lobe’s center. **b,** AP+/PI- cells from *B. schlosseri* testes were analyzed using FACS based on **c,** size (x-axis) and granularity (y-axis). **d-f,** Images of cell populations isolated from *B. schlosseri* testes according to the FACS gates set in **c** (repeated 5 times independently with similar results). Sperm cells from the “SZ” gate **(d, green)**, spermatids from the “STD” gate **(e, orange)**, morphologically undefined cells from the candidate GSC “cGSC” gate **(f, blue)**, and a histogram showing the AP activity gradient in cells from those gates **(g)**. **h-m,** FACS analysis of testis cells throughout development. AP+/PI- cells from *B. schlosseri* testes from primary buds and adult zooids in different developmental stages were analyzed based on size (x-axis) and granularity (y-axis) using FACS. Testis (see red frame) cells from the primary bud of a stage C colony **(h)** and stage D colony **(i)**. Testis cells from the zooid of a stage A colony **(j)**, stage B colony **(k)** and stage C colony **(l)**. Cell fractions in each sorting gate are indicated. **m,** Proportions of the different sorted cell types at each stage from FACS analysis. **n,** Schematic illustration of the *B. schlosseri* weekly developmental cycle. *Apparent PI signal in AP-high cells likely reflects AP-channel spillover.

Studying specific types of stem cells in an organism requires their identification and isolation or enrichment. In *B. schlosseri*, stem cell enrichment was previously achieved using fluorescence-activated cell sorting (FACS) coupled with non-species-specific stem cell features and markers like increased alkaline phosphatase (AP) or aldehyde dehydrogenase (ALDH) activities^11,22^. Laird et al. confirmed stem cell function by transplanting FACS-sorted cells expressing high ALDH activity from donor colonies and showing that these cells successfully contributed to the genetics of recipient colonies^11^. However, a key limitation of this approach is that a high proportion of *Botryllus* cells naturally express ALDH activity, and the cells were enriched from suspensions of entire dissociated colonies and not from a specific cell population or tissue. Therefore, the fact that those cells contributed to either somatic or germline fates suggested that GSCs are separate from mixture of somatic tissue stem cells^11^. This underscores the critical need for strategies to identify and isolate single stem cells in a tissue-specific manner. In fact, Rosental et al. identified hematopoietic stem cells (HSCs) from *B. schlosseri* and demonstrated multilineage differentiation into several blood cell types^22^. However, to date, no protocol has been established for the specific enrichment of GSCs in *Botryllus*, and their biology, aging and fitness-controlling factors remain largely unexplored.

In this set of studies, we found ways to investigate how aging affects reproductive fitness at the cellular level. We developed a method to isolate GSCs and other cell types involved in spermatogenesis in *B. schlosseri*. We characterized their cellular morphology, validated their function in vivo, and tracked their differentiation pathways in vitro and in situ during the organism’s weekly developmental cycle. To identify age-related changes, we generated high-resolution single-cell transcriptomic profiles and compared them between young and old animals, aided by a newly assembled *B. schlosseri* genome. Finally, to shed light on the evolutionary conserved elements of GSC aging from invertebrate chordates to mammals, we analyzed our *Botryllus* single-cell RNA sequencing (scRNA-Seq) dataset in comparison to published mouse and human single-cell germline datasets^23,24^.

## Results

### High-contiguity *B. schlosseri* genome sequencing and assembly

To aid with the transcriptomic analysis in this study we generated, using PacBio platform, a new *B. schlosseri* genome assembly and annotation. Using DNA from the same animal as in our previously published genome^25^ enabled a direct comparison between the earlier *B. schlosseri* genome and the assembly generated in this study. The contiguity of the new assembly, as measured by the N50 value (1,097 kb), was substantially higher than that of our previously published genome^25^ (7 kb; details in Supplementary Table 1). The updated annotation identified 44,727 total genes, including 30,968 predicted genes with introns and 13,759 single-exon genes. With a maximum contig length of 5.27 Mb and a mean contig length of 312,224 bp, this new generated genome assembly provides the sequence continuity needed to study large-scale gene clusters and regulatory regions that were previously obscured by assembly gaps. It also provided complete gene models, as demonstrated by the high percentage of paired-end scRNA-Seq reads that aligned to the new genome (>80%), a rate comparable to mapping benchmarks for the human reference genome^26^.

### Isolation and characterization of GSCs from the testis

*B. schlosseri* is a hermaphrodite species containing both male and female gonads. However, since the female gonad activity in *Botryllus* is limited to the reproductive season, and contain none to only a few oocytes per zooid^14,27^, we focused on the male gonads, which are more abundant and develop weekly. We searched for spermatogonial stem cells (spermatogonia), the type of GSCs that, through spermatogenesis in the testis, give rise to mature sperm (Fig. 1a). Using FACS, we sorted 3 different cell populations based on size and granularity of cells expressing increased AP activity (Fig. 1b-g). AP activity has been used to select for GSCs in mammals^28^ and we hypothesized that it may be a characteristic of stem cells in the gonads of protochordates. Increased ALDH activity was also used previously to select for stem cells in mammals^29,30^ and marine invertebrates^31^, including *B. schlosseri*^11^, and FACS analysis of cells expressing high ALDH activity in the testis of *Botryllus* yielded the same cell populations as AP (Extended Data Fig. 1). AP and ALDH were used as independent stem-cell-associated markers to validate our enrichment strategy, as both have previously been used to enrich for stem cells in *B. schlosseri*^11,22^, although not specifically from the gonads.

Using this sorting strategy, we identified the sperm and spermatid cell populations (Fig. 1d, e) but could not define the cells in the other population (Fig. 1f). However, since the sperm and spermatid populations were clearly distinguishable, and since the AP activity level in the morphologically undefined cell population was much higher (Fig. 1g), we hypothesized that the latter contains the spermatogonial stem cells and named it as candidate germline stem cell (cGSC) population. To identify the GSCs and further characterize the different cell types in *Botryllus* testis, we index sorted 3,264 cells, using FACS, from the sperm, spermatid and cGSC populations into 96-well plates for full-length scRNA-Seq using Smart-seq3^32^. Index sorting was used because it records the fluorescence intensities for each sorted cell, allowing for direct correlation between its transcriptome and its exact location on the FACS plot. To include all possible cell types involved in spermatogenesis, the cells for the scRNA-Seq were isolated from testes at different time points along the *Botryllus* developmental cycle including adult zooids in stage A and primary buds and adult zooids in stage C. The developmental stages are illustrated in Fig. 1n and were defined according to Lauzon et al.^33^ (as explained in Supplementary Table 2). Alignment of the scRNA-Seq and FACS data showed that the transcriptome-based cell clusters (Fig. 2a) clearly distinguished the different sorted cell populations (cGSC, spermatid and sperm) (Fig. 2b). We annotated each cluster based on gene profiles of known cell-type-specific markers (Fig. 2c,d, Supplementary Table 3). While all cells annotated as spermatogonia and 85% of cells annotated as spermatocytes were sorted from the cGSC gate, this gate also contained smaller proportions of support cells (3.70%), differentiating cells (8.50%), spermatids (11.25%), and sperm (3.12%). Although we expected scRNA-seq to resolve developmental subpopulations within major cell types, such as primary versus secondary spermatocytes and round versus elongated spermatids, further high-confidence annotation was limited due to the large number of *Botryllus*-specific markers lacking functional annotation (Fig. 2d). Based on the scRNA-Seq analyses, *DDX4,* (*VASA*)^23,34,35^ and *PIWIL1*^35–37^ were used as representative markers for spermatogonia; *SYCP3*^23,35,38^ and *H1F0*^39,40^ were markers for spermatocytes, *PSME3*^41–43^, *UBE2R2*^41,42,44^ and *SAXO4*^41,42,45^ (formerly *PPP1R32*) for spermatids and *PRM1*^46^, *SPA17*^47^ and *TSSK1B*^48^ for sperm (Fig. 2e-h, Supplementary Table 4). Cluster 16 did not express specific markers for cells involved in spermatogenesis but expressed high levels of *ACTIN* transcripts. We hypothesize that this cluster may contain *ACTIN* RNAs from myocytes and other cells expressing cytoskeletal actins for their own functions. Most of the highly expressed genes of Cluster 6 were not annotated and those that were annotated could be correlated with either spermatocytes, spermatids or sperm. Therefore, we hypothesize that this cluster contains actively differentiating cells from spermatocytes to spermatids and sperm. Importantly, Cluster 10, which contains cells from the cGSC FACS gate, exhibited strong expression of spermatogonia-associated genes such as *DDX4*^23,34,35^ and *PIWIL1*^35–37^, germline markers like *TDRD7*^49^, along with genes linked to proliferating cells such as *PCNA*^50^. This suggests that we successfully identified GSC population in *B. schlosseri* testis and developed a reproducible FACS-based method to isolate them, enabling us to further investigate their capacity to engraft and differentiate in vivo, as well as to explore their aging dynamics.

**Fig. 2:**
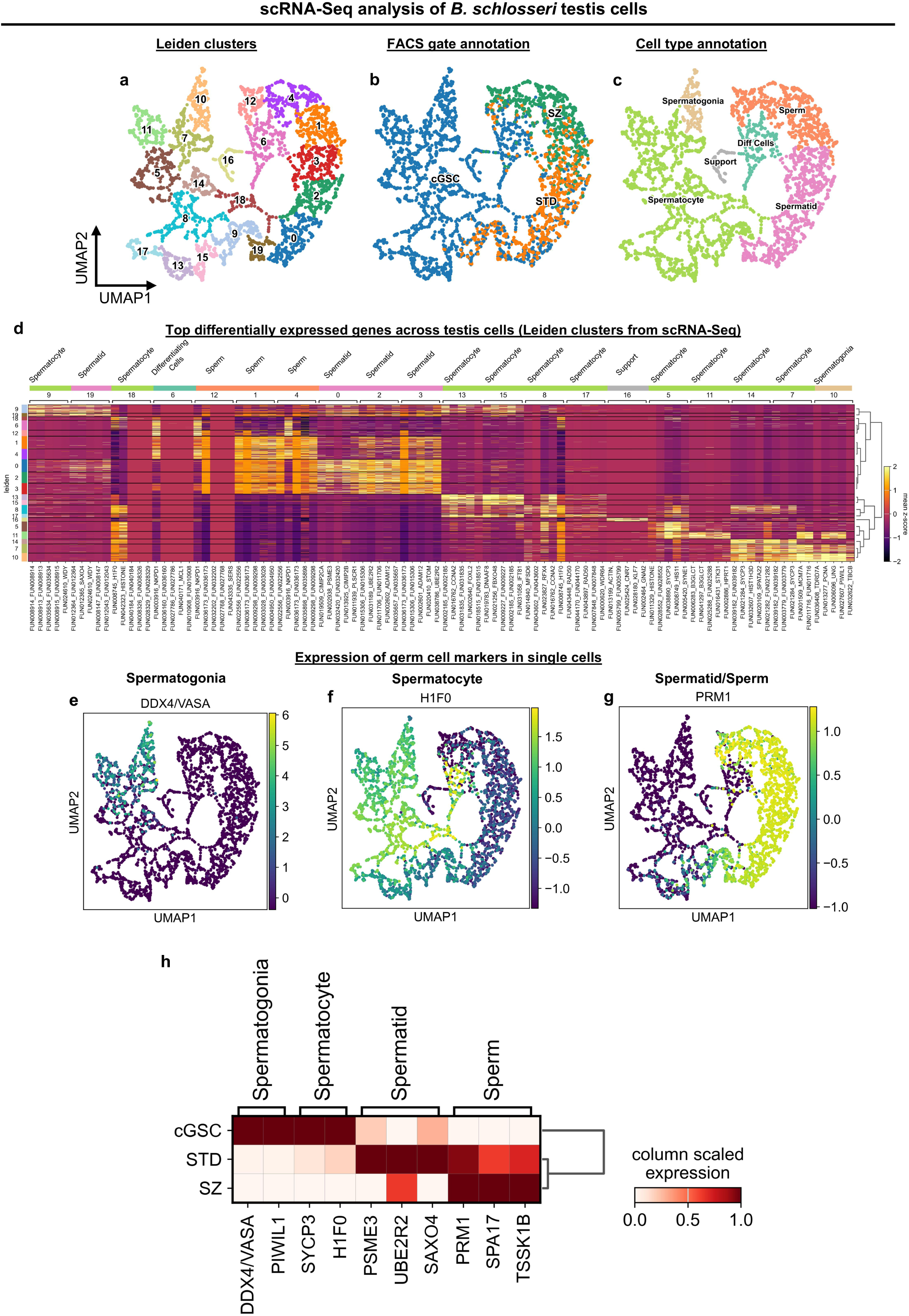
A single-cell atlas of *B. schlosseri* spermatogenic cells. **a-c,** scRNA-Seq of index-sorted AP+/PI- testes cells using Smart-seq3. **a,** UMAP plot showing the numbered Leiden clusters. **b,** UMAP plot with annotated clusters based on the FACS gate by which the cells were sorted. **c,** UMAP plot with annotated cell type identities based on expressed transcripts of known genes **(d)**. Related to Supplementary Table 3. **e-h,** scRNA-Seq expression analysis of representative germ cell markers. UMAP expression plots of representative gene markers for different cell types that were used for in situ expression analysis (HCR; see Fig. 5). **e,** *DDX4* – spermatogonia. **f,** *H1F0* - spermatocytes. **g,** *PRM1* – spermatids and sperm. **h,** Matrix plot showing marker gene expression for transcriptomically defined clusters from scRNA-Seq and the FACS gate they were sorted from.

### Identifying primordial germ cells in the cell islands niche

The *B. schlosseri* cell islands (Extended Data Fig. 2a) form a putative niche for both germ and somatic stem cells^17^. Primordial germ cells (PGCs) are early germline precursors that give rise to GSCs (like spermatogonia) during development, which then maintain and generate the germline^51^. To evaluate the presence of PGCs and GSCs in that niche, we index-sorted cells with high AP activity from dissected cell islands isolated from adult Stage A zooids (Extended Data Fig. 2b). A total of 384 index-sorted cells were analyzed by scRNA-Seq (Extended Data Fig. 2c). Multigene analysis showed that Cluster 10 expressed prominent markers of PGCs such as *NANOS3*^52^ and *AGO1*^53^ (Extended Data Fig. 2d, e, Supplementary Table 5). This indicates that, at the transcriptomic level, the cell islands may contain PGCs. It is consistent with a previous transplantation study in which the donor genotype was detected in the recipient testis following whole cell islands transplantation, suggesting the presence of gonad-forming stem cells in the cell islands niche^17^. Furthermore, representative assay using in situ hybridization chain reaction (HCR) for *AGO1* showed high expression in the testis, suggesting its high specificity to the germline niche in *Botryllus* (Extended Data Fig. 3). Based on gene expression profiles (Extended Data Fig. 2g, Supplementary Table 5), the other clusters were found to include gonadial cells, muscle cells, epithelial cells and hemocytes (Extended Data Fig. 2f). While the cell islands were previously suggested to contain heterogenous cell types including both somatic and stem cells^17,54^, their small size, embedded location, and proximity to surrounding tissues, made precise dissection challenging. As a result, cells from adjacent tissues, including testis, may have been inadvertently collected during dissection, yielding a heterogenous single-cell dataset despite the relatively small number of captured cells.

### GSC engraftment and differentiation in vivo

A fundamental trait of stem cells is their ability to give rise to different cell types while simultaneously maintaining a self-renewing stem cell pool. In vivo transplantation and tracing of cells in non-model organisms is challenging due to the lack of ubiquitous reporter lines. Therefore, to investigate the in vivo capacity of the cGSC population to engraft and differentiate into sperm in *B. schlosseri*, we developed a novel method combining transplantation and scRNA-Seq. This approach involved transplantation of dye-labeled donor cGSCs into compatible syngeneic recipients (to avoid histocompatibility-related rejection^55–57^) followed by index sorting and scRNA-Seq. PGCs begin to accumulate in the secondary buds^16^ (Extended Data Fig. 4), where they subsequently give rise to the developing GSC population (spermatogonia) during gonadal development. However, due to their small size, we could not dissect their gonadal niche. Therefore, to enrich for PGCs and GSCs from both the buds and zooids, for the transplantation assay, we applied the FACS gates that we developed from pure testes to a cell-suspension produced from whole animals at stage B. From this suspension, we sorted cells with high ALDH activity. ALDH was selected to incorporate another marker that was previously used for stem cell identification in *B. schlosseri*^11^, and it showed similar staining patterns compared to AP as part of our FACS gating strategy (Extended Data Fig. 1). To validate that the GSC sorting gate from whole animals enriches for GSCs, we performed bulk RNA-seq on the GSC-gated population and the total live-cell population from 7 independent biological replicates derived from 4 distinct *Botryllus* colonies bearing different genotypes. Differential expression analysis demonstrated that the GSC-gated population expresses higher levels of prominent germline markers, including *DDX4*, *PIWIL1*, *PIWIL2*, and *NANOS3*, confirming that this sorting strategy from whole animals enriches for GSCs (Extended Data Fig. 5, Supplementary Table 6). ALDH+ cells were sorted based on the cGSC gate (Fig. 3a), labeled with a fluorescent membrane dye (DiD), and transplanted into recipient animals at stage B through injection into their vasculature. Seven days post transplantation (one developmental cycle), we used FACS to analyze cells from the gated “Testes” population (combining cGSCs, spermatids, and sperm) from the whole animals, that reached to stage B again and captured 1,726 cells from the DiD+ gate for scRNA-Seq (Fig. 3b). Control animals were subclones of the same genotype and developmental stage but not transplanted. Cells from these control animals, which exhibit natural autofluorescence in the DiD+ gate, were sorted for sequencing and labeled as “control”. Cells from the transplanted animals which potentially contained gametes from engrafted GSCs, in addition to autofluorescent cells in the DiD+ gate, were sorted for sequencing and labeled as “transplanted” (Fig. 3b). Clustering analysis of the DiD+ cells emphasized Leidens 15 and 10, which highly expressed the germ cell markers previously investigated in this study (*DDX4*, *PIWIL1*, *SYCP3*, *H1F0*, *PSME3*, *UBE2R2*, *SAXO4*, *PRM1*, *SPA17*, and *TSSK1B*) (Fig. 3c,d). Sub-clustering of these clusters enabled the identification of clusters containing spermatogonia and spermatocytes (Leidens 3, 0, 7, 2) and those containing spermatids and sperm (Leidens 4, 1) (Fig. 3e,f). We then determined the origin of the annotated cell populations and found that ∼60% of the spermatogonia and spermatocytes and ∼80% of the spermatids and sperm were DiD+ (Fig. 3g,h), indicating that they originated from the GSC-transplanted animals. These results demonstrate that the transplanted GSCs not only remained viable and contributed to the formation of the recipient testis, but also that a substantial proportion of them differentiated into spermatocytes, spermatids, and mature sperm. To further validate the fate of transplanted GSCs, we performed two complementary experiments. First, we tested whether GSCs preferentially localize to the recipient’s testes compared with a control cell population. DiD-labeled GSCs were transplanted into one group of recipient animals (n=3), while DiD-labeled pigment cells^22^ (as a control) were transplanted into a separate group of recipients (n=2). Seven days post transplantation, cell suspensions were prepared from recipient testes and analyzed by flow cytometry. The proportion of DiD+ cells detected in the testes of recipients transplanted with GSCs was higher than that observed in recipients transplanted with pigment cells, indicating that GSCs localized to the testes while pigment cells didn’t (Fig. 4a-c). Moreover, we performed a transplantation experiment in which DiD-labeled GSCs and DiO-labeled sperm cells (as a control) were co-transplanted into the same recipients (Fig. 4d) and analyzed by confocal microscopy 7 days post transplantation. Because cell islands contain abundant number of phagocytes that may capture transplanted cells, we first examined cell islands as a control tissue to confirm successful transplantation. Both DiD+ and DiO+ cells were detected in the cell islands, demonstrating that both labeled cell populations were successfully introduced into the recipients. In contrast, analysis of dissected recipient testes revealed DiD+ cells but no detectable DiO+ cells. Notably, some DiD+ cells exhibited sperm-like morphology, suggesting that transplanted GSCs not only localized to the testes but also differentiated into sperm within 7 days following transplantation (Fig. 4e). Together, these results strongly suggest that we not only successfully isolated a GSC-enriched cell population but also revealed that these cells were functionally viable to home to the gonadal niche of the recipients, engraft, and undergo differentiation into spermatocytes, spermatids and sperm in host animals.

**Fig. 3:**
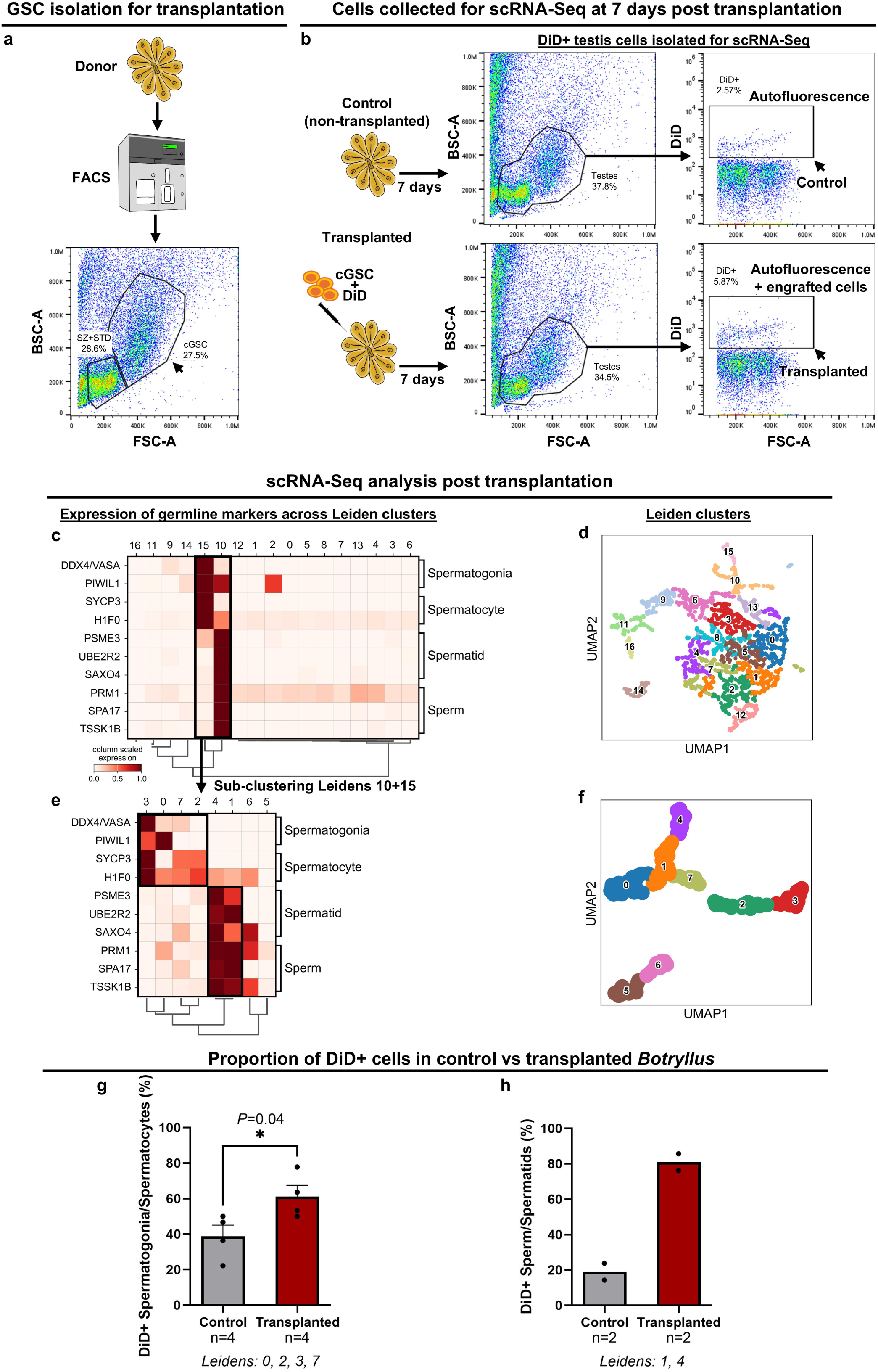
Engraftment and differentiation of transplanted candidate GSCs revealed by in vivo tracing and scRNA-Seq. **a,** ALDH+ cGSC cells (black arrow) were sorted, based on our gating strategy, from a whole animal using FACS and transplanted into recipient colonies. The cells from the cGSC gate were labeled with DiD prior to transplantation to recipients (n=3) while control animals were not transplanted (n=2). **b,** Seven days post transplantation (after one regeneration cycle and takeover) DiD+ cells (black arrows) from the Testes gates (cGSC+STD+SZ) applied on cells coming from a whole-animal cell suspension were index-sorted for scRNA-Seq, using Smart-seq3, from transplanted and non-transplanted control animals, which were at the same stage and had visible testes. In the control animals, the cells from the DiD+ gate displayed natural autofluorescence, while in the transplanted animals, this gate contained both naturally autofluorescent cells and engrafted cells. **c,** Matrix plot of germline marker expression (*DDX4, PIWIL1, SYCP3, H1F0, PSME3*, *UBE2R2*, *SAXO4*, *PRM1*, *SPA17*, *TSSK1B*) for each Leiden cluster showing the highest expression in Clusters 10 and 15. **d,** UMAP plot showing the numbered Leiden clusters. **e,** Matrix plot of germline marker expression after sub-clustering of Clusters 10 and 15. **f,** UMAP plot showing the numbered Leiden clusters in the sub-clustered data. **(g-h)** Quantification of DiD+ cells in control vs transplanted animals. DiD+ cells were more abundant in transplanted animals than in controls in both spermatogonia/spermatocytes **(g)** and spermatids/sperm **(h)** (each point represents one Leiden cluster assigned to the indicated cell type; n represents the number of Leiden clusters analyzed; statistical significance: two-tailed unpaired t test with Welch’s correction is indicated where applicable; data are presented as mean values ± SEM, where shown). Additional details of the ALDH+ GSC enrichment and associated marker enrichment analysis are provided in Extended Data Fig. 5.

**Fig. 4:**
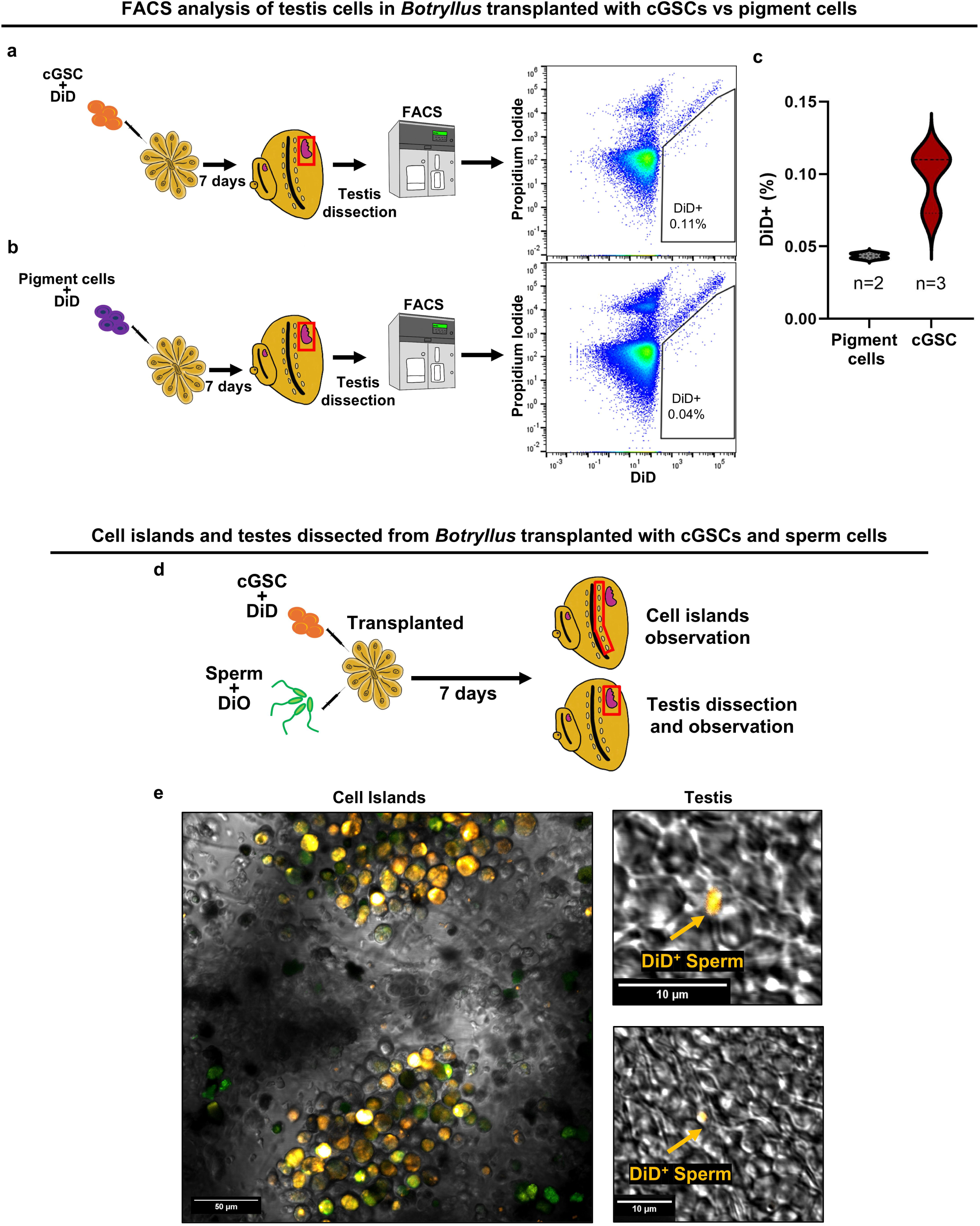
Engraftment and differentiation of transplanted candidate GSCs revealed by FACS analysis and confocal imaging. **(a–c)** FACS-based detection of donor-derived cells in recipient testes after transplantation. DiD-labeled cGSCs **(a)** or DiD-labeled pigment cells as control **(b)** were transplanted into recipient compatible subclones of the same colony. cGSC recipients (n=3) and pigment-cell recipients (n=2). Seven days post transplantation, testes (red frame) were dissected, dissociated, and analyzed by flow cytometry. Representative FACS plots show the DiD+ gate for cGSC-transplanted and pigment-cell–transplanted recipients. **(c)** Quantification shows a higher percentage of DiD+ cells in dissociated testes from cGSC-transplanted versus pigment-transplanted recipients. **(d)** In a separate experiment, DiD-labeled cGSCs (orange) and DiO-labeled sperm cells as control (green) were co-transplanted into the same recipients. Tissues for confocal observation are labeled with red frame. **(e)** Seven days post transplantation, cell islands were imaged to confirm delivery of both labeled cell types to the recipients (as indicated by phagocytes engulfing both DiD+ and DiO+ cells). Confocal imaging of dissected testes from the same recipients revealed DiD+ cells with sperm morphology (arrows), whereas DiO+ sperm were not detected in the testes, suggesting that transplanted cGSCs, but not mature sperm, can engraft within the recipient’s testis and differentiate into sperm.

### Cellular-level spatiotemporal dynamics of gonad development

*B. schlosseri* colonies undergo a complete whole-body regeneration cycle every week^16,33^. As a crucial part of this process, PGCs must migrate from resorbed zooids to the developing buds to establish the gonads of the next generation^14,16^. To follow testis development at the cellular level, we dissected testes from primary buds at stages C and D, and from adult zooids at stages A, B and C. Testis dissection from primary buds at earlier stages and from secondary buds was not feasible due to their small size. Using FACS, we analyzed the fraction of the cGSC, spermatid and sperm cell populations at each developmental stage (Fig. 1h-l). While the cGSC population was consistently present in testes across all tested parts and stages, spermatids first appeared in the primary bud at stage D, and sperm first observed in the adult zooid at stage A. The proportions of each population along the developmental cycle show that GSCs gradually differentiate into spermatids and sperm, with the highest fraction of sperm cells observed in the testes of adult zooids at stage C (Fig. 1l). These results reveal that at stage C, while the testis of the primary bud contains mostly GSCs, in the adult zooid within the same animal, most GSCs have already differentiated into spermatids and sperm. This is a remarkable coexistence of 2 asexual generations within the same individual having testis at different developmental stages. Since we couldn’t dissect testis from secondary buds, we used histology and 3D reconstructions to identify the earliest developmental stage at which gonads begin to form (Extended Data Fig. 4). A few PGCs were found in the gonadal niche of the secondary bud at stage B (Extended Data Fig. 4a, b), and more organized PGCs along with early developing oocytes were observed in the secondary bud at stage C (Extended Data Fig. 4c-f).

To further investigate gonad development and validate our scRNA-Seq results, we examined the expression pattern of 3 marker genes from our scRNA-Seq analysis using HCR on whole-mount animals along the weekly developmental cycle. The markers chosen were *DDX4* (*VASA*; for GSCs^34^), *H1F0* (for spermatocytes^39,40^) and *PRM1* (for spermatids/sperm^46^) (Fig. 5, Extended Data Fig. 6). *DDX4* expression began in PGCs in the secondary bud at stage B (Fig. 5d) and in oocytes in the secondary bud at stage C (Fig. 5e). Its expression was clearly visible in the developing testis in primary buds from stages B to D (Fig. 5d-f), but disappeared once the primary buds transitioned to stage A zooids (Fig. 5c). *H1F0* expression was testis-specific, beginning in the primary bud at stage C (Fig. 5j). It remained strong at stage D (Fig. 5k) but began to fade in the adult zooids at stages A and B. This fading coincided with the differentiation of spermatogonia and spermatocytes into spermatids and sperm, as demonstrated by the increasing expression of *PRM1* in the center of the testis lobes (Fig. 5h, i). Strikingly, aligned with our FACS data of testes from primary buds and adult zooids (Fig. 1h, l), the in situ gene expression profiles of *H1F0* and *PRM1* at stage C also demonstrated the coexistence of testes from 2 different asexual generations within the same animal (Fig. 5j).

**Fig. 5:**
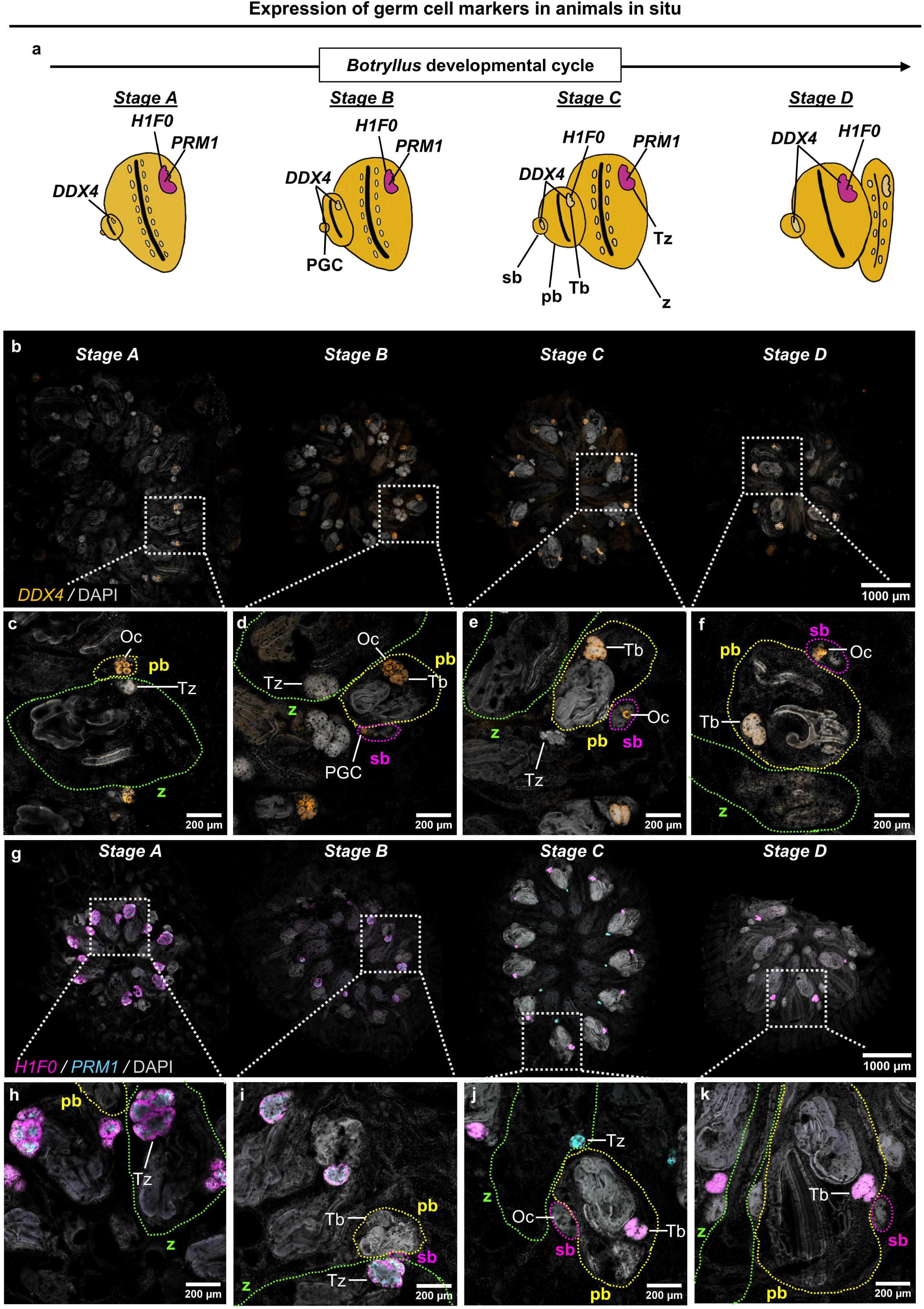
Spatiotemporal expression of germ cell markers during gonadal development. **a,** Schematic summary of germ cell marker expression across developmental stages of the *Botryllus* weekly cycle. Testes in the zooids and buds are stained magenta and light yellow, respectively. **b-k,** Germ cell marker expression during the *B. schlosseri* developmental cycle. **b,** HCR images showing *DDX4* (orange) expression which begins in the developing gonad of the secondary bud in stage B. The gray frames in **b** mark the close-ups shown in **c-f**. Close-up images of a colony in stage A **(c)**, stage B **(d)**, stage C **(e)** and stage D **(f)**. **g,** HCR images showing *H1F0* (magenta) and *PRM1* (cyan) expression which begins in the developing gonad of the primary bud in stage C and the zooid in stage A, respectively. The gray frames in **g** mark the close-ups shown in **h-k**. Close-up images of a colony in stage A **(h)**, stage B **(i)**, stage C **(j)** and stage D **(k)**. Specimens are counterstained with DAPI (gray). The zooid (z), primary bud (pb) and secondary bud (sb) are outlined with green, yellow and magenta frames, respectively. The primordial germ cells (PGC), developing oocytes (Oc), zooid’s testis (Tz) and primary bud’s testis (Tb) are marked. Additional close-up images are shown in Extended Data Fig. 6. The experiment was repeated 3 times independently with similar results.

### Age-associated molecular changes in GSCs across species

Reproductive capacity declines with age in many animals, and reproductive aging is associated with conserved changes in GSC maintenance, proliferation, and differentiation capacity across species^5,23,24,58,59^. These changes can affect fertility and contribute to age-related declines in reproductive function. Using our newly developed strategy to isolate GSCs in a protochordate, we investigated their role in gonadal aging at the single-cell-transcriptomic level and compared these findings to mice and humans. Using FACS, we index-sorted and analyzed 1,968 single cells with high ALDH activity from the testes of young (6-months old) and old (163-months old) *Botryllus* colonies in stage A. Cell suspensions were made from testes collected from multiple zooids across several systems within the same colony. The cells were sorted from both the GSC population which is enriched with spermatogonia and from the STD+SZ population containing spermatids and sperm (Fig. 6e). Clustering analysis demonstrated clear separation between cells derived from young and old colonies (Fig. 6a-c). Using MAST differential expression analysis^60,61^, we identified distinct marker genes for each cell type and age group. Remarkably, based on gene expression of the top differentially expressed genes, we observed a higher variation between young and old GSCs compared to young and old spermatids/sperm (Fig. 6d, Supplementary Table 7). These results show that age-associated gene expression differences are more evident in GSCs than in spermatids/sperm. However, whether these molecular differences contribute to the maintenance of reproductive function in aged colonies or instead reflect reduced reproductive capacity in aged colonies are yet to be determined. To further investigate the germline single-cell-transcriptomic patterns in young and old *Botryllus* colonies, we analyzed the expression level of the main gene markers studied throughout this project. The expression of markers associated with spermatogonia and spermatocytes (*DDX4*, *PIWIL1*, *SYCP3*, *H1F0*) was higher in young compared to old colonies, while markers associated with spermatids and sperm (*PSME3*, *UBE2R2*, *SAXO4*, *PRM1*, *SPA17*, *TSSK1B*) showed the opposite pattern (Fig. 6f, Extended Data Fig. 7a,b). A comparative meta-analysis of scRNA-Seq dataset from mouse testes^24^ showed a similar expression pattern for the same set of genes among cells from young (2-months old) and old (24-months old) animals (Fig. 6g, Extended Data Fig. 7c). However, the expression pattern of those genes was slightly different in humans. The meta-analysis of scRNA-Seq data of human testis cells^23^ showed higher expression in younger men for *PIWIL1* and *PRM1*, little difference for *DDX4*, and lower expression levels for *SYCP3*, *H1F0*, *PSME3*, *UBE2R2*, *SAXO4*, *SPA17* and *TSSK1B* (Fig. 6h, Extended Data Fig. 7d). Still, it is outstanding that even with the high variation between human samples of the same age (Extended Data Fig. 7d), which may be attributed to factors like elevated body weight, smoking^23^ or simply diverse genetics, the overall trend of higher expression of the GSC marker *PIWIL1* and lower expression of most tested spermatid/sperm markers (with the exception of *PRM1*) in younger compared with older samples were consistent with the patterns observed in mice and *Botryllus*. This strongly suggests the existence of a conserved reproductive aging mechanism that has been maintained across hundreds of millions of years of evolution since the divergence of protochordates and mammals.

**Fig. 6:**
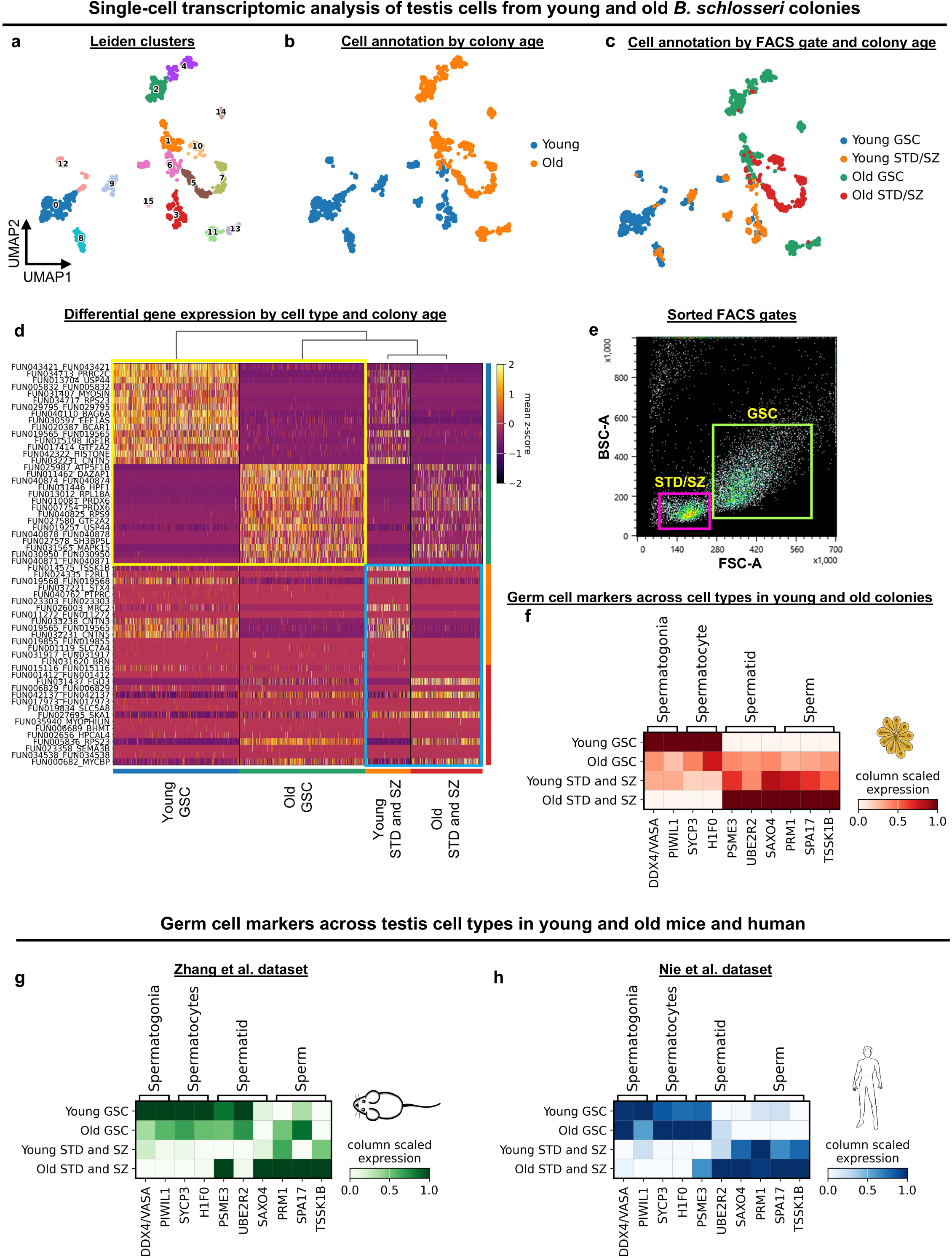
Germline stem cell transcriptomic alterations influence reproductive aging and reveal conserved marker profiles across chordates. **a-e,** scRNA-Seq of index-sorted testes cells from young and old *B. schlosseri* colonies using Smart-seq3. ALDH+/PI- cells from the GSC and STD+SZ gates were sorted using FACS based on morphological parameters. **a,** UMAP plot showing the Leiden clusters with cluster numbers. **b,** UMAP plot with annotated clusters based on the age of the animal from which the cells were sorted. **c,** UMAP plot with annotated clusters based on the FACS gate and the age of the animal from which the cells were sorted reveals distinct cell clusters of young (blue) and old (green) GSCs, as well as young (orange) and old (red) sperm and spermatids. **d,** A heatmap plot displaying the top 15 differentially expressed genes (based on Supplementary Table 7) in young and old GSCs (yellow frame) and young and old sperm and spermatids (cyan frame) further illustrates the higher aging-related transcriptomic variation within the GSC population compared to spermatids and sperm. **e,** FACS gates used for sorting cell populations for scRNA-Seq. **f-h,** conserved germline stem cell marker profiles underlie reproductive aging across chordate evolution. Matrix plot showing marker gene expression for defined cell types (*DDX4*, *PIWIL1*, *SYCP3*, *H1F0*, *PSME3*, *UBE2R2*, *SAXO4*, *PRM1*, *SPA17*, *TSSK1B* – details in Supplementary Table 4) from scRNA-Seq in cells according to the animal’s age and cell type in *B. schlosseri* (f), *M. musculus* (g) (10X Genomics data from Zhang et al., 2023) and *H. sapiens* (h) (10X Genomics data from Nie et al., 2022). Related to Supplementary Tables 8-9. Expanded matrix plots (including biological replicates for mouse and human) and detailed gene marker comparisons between young and old *Bortyllus* are shown in Extended Data Fig. 7.

## Discussion

Reproductive aging, a universal phenomenon impacting fertility across the animal kingdom, remains a significant biological puzzle. Previous single-cell studies reported varying degrees of age-related transcriptomic differences in GSCs, ranging from distinct^24,62^ to modest^23^. However, these studies used droplet-based scRNA-Seq platforms on unsorted testis cell suspensions, which likely included transcriptional noise from unrelated cell types present in the heterogeneous cell samples. To overcome this limitation, we employed a plate-based method combined with index sorting to analyze purified testis cell populations, enabling high-resolution analysis of age-related changes within defined cell types.

With the aim to characterize the different cell types involved in *B. schlosseri* spermatogenesis, we combined 2 previous approaches that used FACS and AP/ALDH activity indicators as non-species-specific stem cell markers^11,22,31^. We employed it on cell suspension from dissected germline niches (testis and cell islands) followed by scRNA-Seq analysis, in vivo transplantation and in situ tracing for validation of stem cell differentiation. This is a generalized method for characterizing tissue-specific stem cells in non-classical model organisms, which often lack transgenic tools and gene-specific monoclonal antibodies. Using this approach, and based on known prominent gene markers, we report here the first high-resolution identification and isolation of the different cell types involved in *Botryllus* spermatogenesis including spermatogonia, spermatocytes, spermatids and mature sperm. While the cell islands in *B. schlosseri* were previously suggested to include GSCs^17^, the presence of diverse cell types within this niche made their definitive identification elusive. Our results here provide a cellular-level analysis that reveals the transcriptomic signature of PGCs, thus supporting the claim that the cell islands might be a GSC niche^17^. Furthermore, by tracking germline markers, we precisely mapped the weekly developmental cycle of the *B. schlosseri* testis at the cellular level. Our results suggest that this process begins with GSC accumulation in the secondary bud at Stage B, progresses through differentiation, and ends two weeks later with sperm release from the testis of the adult zooid at Stage C. The expression pattern of the sperm/spermatid-related marker, *PRM1*, aligned with the sperm release temporal pattern that was previously reported for this organism^63^. Accordingly, we suggest that PGCs either initially develop in the cell islands and then migrate to the secondary bud, where they initiate testis formation and differentiate into spermatogonia, or that they originate in the secondary buds and then transit through the cell islands to the developing testes.

The strategy we have developed to enrich for GSCs and other germline cell types allowed us to investigate their aging dynamics at the single-cell level. A core strength of *Botryllus* as a model lies in its unique life history, in which the aging of a colony is suggested to be dictated by the aging of its stem cells, which self-renew and migrate each week across asexual generations^12^. The remarkable 163-months old colony that was used in this study represents approximately 700 asexual generations of persistent stem cell activity. Although aged stem cells may be prone to higher mutation rates^5^, age-associated reproductive decline may not arise solely from accumulated genetic alterations. Other potential factors include epigenetic regulation, transposable-element insertion into regulatory regions, or simply deterioration of the gonadal niche, which could affect GSC function and gene expression without necessarily altering the DNA sequence.

Our study reveals a strikingly distinct aging pattern among different cell types within the *Botryllus* male germline. GSCs exhibit substantial age-related transcriptional changes, while their differentiated gamete progeny maintains relatively stable transcriptomic profiles. This central finding suggests that molecular differences in GSCs, rather than in differentiated germ cells, might either maintain reproductive capacity in aged colonies or mark a decline in reproductive success. Specifically, we found a significant age-related decline in the expression of the prominent GSC markers *DDX4* and *PIWIL1*. These genes are essential for GSC function and fertility in many organisms^34,36,64,65^, including protochordates^66–68^ and thus may constitute a conserved mechanism for maintaining reproductive fitness across species.

Our comparative analysis revealed similar expression patterns of spermatogenesis-related gene markers in both *Botryllus* and mice between young and old testes cells but a slightly different pattern when comparing cells from young and old human testes. The substantial variation between human samples, potentially due to lifestyle factors like smoking and increased body weight^23,69^, might have contributed to differences in the gene expression pattern compared to *Botryllus* and mice. Interestingly, despite this high variability observed between human samples which can be attributed to genetic or health-related factors, the consistent age-related decline in *PIWIL1* expression in human germline cells mirrored the pattern observed in *B. schlosseri* and mouse. These results support our central hypothesis that reproductive fitness and its underlying molecular and physiological factors can be reflected by molecular changes in GSCs and are likely conserved across evolution, from simpler to more complex organisms.

This is not the first demonstration of a GSC-related mechanism showing a conserved pattern among protochordates and mammals. For instance, the GSC competition phenomenon observed in *Botryllus*^9,10^ prompted us to search for a similar mechanism in mammals^15^. We have previously shown that the germline in mice is derived from only 4 cells at the epiblast stage^15^. Transplantation of color-coded embryonic stem cells into uncolored blastocysts produced tetrachimeric mice with cells of all colors in most of their tissues, including the fetal and neonatal testes. However, during the reproductive phase, most GSCs underwent apoptosis, and only the surviving cells contributed to the mature gonad^15^. These remarkable findings suggest the existence of GSC competition across protochordates^8–11^ and mammals^15^, but the underlying mechanisms that determine GSC fitness remain largely unclear.

Investigating the driving factors of ‘winner’ GSCs is highly important since GSCs are the only stem cells capable of passing genes to future generations, making them fundamental units of natural selection^8,11^. Therefore, the present study, which reports the first isolation of GSCs in this unique protochordate model is a foundational step towards identifying the molecular and cellular factors that govern GSC competition. Moreover, the evolutionary position of protochordates, as the likely closest living invertebrate relatives to vertebrates^70^, makes the present study of their GSCs highly valuable from a multidisciplinary perspective, as insights from these organisms may be further applied to more complex organisms, including mammals. From an ecological standpoint, understanding how GSCs function can pave the way for innovative strategies to conserve endangered species by addressing the impacts of environmental stress on their GSC fitness and on their reproductive health. On a broader scale, in the fields of regenerative medicine and reproductive aging, the fact that basic biological mechanisms, such as the aging of GSCs explored in this study, and GSC competition, seem to be conserved across protochordates and mammals, suggest that our findings and the techniques established to isolate GSCs in a non-classical model organism like *B. schlosseri* can potentially be harnessed in the future to develop therapies to improve human reproductive health.

## Methods

### Animal husbandry

*B. schlosseri* mariculture procedures have been previously described^71^. Animals were collected from the marina in Monterey, California and transferred to the mariculture facility at Hopkins Marine Station. The animals were tied to 3 × 5-cm glass slides while another empty glass slide (to collect hatches) was placed in a slide rack in front of the slide with the tied animal. The slide rack was placed in an aquarium with filtered seawater, and within a few days the sexually reproduced larvae hatched, swam to the empty slide, settled and metamorphosed into the adult body plan. Single hatched animals were then transferred to individual slides, allowing them to form colonies that were used in the experiments. Each colony originated from a single sexually produced larva and expanded through weekly blastogenic cycles (asexual budding), generating genetically identical zooids and buds. During each cycle, adult zooids are resorbed and replaced by developing buds. Therefore, colony age refers to the time since larval settlement rather than the lifespan of an individual zooid.

### Tissue dissociation

To avoid food contaminants coming from the digestive system in our experiments, the animals were isolated without food for 24 hours prior to dissection. Insulin syringes were used to dissect testes and cell islands from laboratory grown *B. schlosseri* colonies. The dissection procedure was performed under a light stereomicroscope, and dissected tissues were collected into 96-well clear flat bottom plates filled with a serum-free “*Botryllus* medium” (Medium 199 Hanks’ Balanced Salts with phenol red and 3.3x calcium/magnesium-free HBSS) and placed on ice until used for further analysis. The testis is a cloudy-looking white tissue located on one or both sides of the developing buds and adult zooids. The cell islands are cell aggregates aligned laterally on both sides of the endostyle niche^17,18,22^ (Extended Data Fig. 2a). The initial incisions were made through the tunic and mantle layers. The branchial sac and digestive system were then gently displaced or removed to expose the tissue of interest. To minimize carryover of surrounding tissue and debris, isolated samples were moved away from the dissection site before transferring to 96-well plates.

### FACS analysis

Cell suspensions from dissected testes (n=15) and cell islands (n=30) were generated by mechanical dissociation using insulin syringe plunger and filtered through a 70-μm mesh followed by a 35-μm mesh into a 5 ml tube containing serum-free *Botryllus* medium. No enzymatic dissociation was carried out. Cells were centrifuged at a speed of 500 G for 5 min, supernatant was discarded, and cells were resuspended in 1 ml of fresh *Botryllus* medium. To enrich for stem cells, 1 μl of Alkaline Phosphatase Live Stain (Thermo Fisher) or ALDH (ALDEFLUOR Kit, Stemcell Technologies) was added. Cells were incubated for 30 min on ice under dark conditions, and 1 μl of Propidium Iodide (PI) (1µg/ml) was added before flow analysis in order to separate live from dead cells. After gating for live PI negative cells, we analyzed the cells using forward (FSC) and back-scatter (BSC), and fluorescence from the AP/ALDH staining using a Sony MA900 FACS instrument. Flow cytometry data were analyzed using Flowjo v10.10 Software (BD Life Sciences). For morphological observations, the sorted cells were collected in 5 ml tubes containing 1 ml *Botryllus* medium and centrifuged at a speed of 500 G for 5 min. The supernatant was discarded and the remaining medium (∼30 µl) was spined down, gently mixed and transferred to µ-slide 18 well glass bottom chamber (Ibidi). The chamber was placed on ice for 20 min to allow cells to settle down followed by imaging using Zeiss LSM700 confocal microscope.

### In vivo transplantation of GSCs

For the first transplantation experiment we used reproductive (visible testes) syngeneic animals (compatible subclones of the same genotype). Cell suspension from a donor whole animal (stage B) was generated using mechanical dissociation as described above. Cells with high ALDH activity from the cGSC FACS population were sorted, centrifuged, resuspended in 500 µl of *Botryllus* medium and labeled with a 5 µl of Vybrant DiD Cell-Labeling Solution (Invitrogen). The cells were incubated for 1 h in room temperature under dark conditions, washed twice with *Botryllus* medium (first with 2% FCS and then with serum-free medium) and resuspended in 30 μl of fresh serum-free medium before injection. For injection, glass needles were prepared using a P-1000 micropipette puller (Sutter Instruments). Needles were loaded with 5 μl of cell suspension (equivalent to ∼30,000 cells) and injected using manual micromanipulator (World Precision Instruments) under a stereoscope into the proximal ampulla of recipient animals, also in stage B (n=3). Non-transplanted, stage B, subcloned animals of the same genotype (n=2) served as a control. Seven days after transplantation, following one regeneration cycle and takeover, we used our FACS gating strategy to sort GSCs, spermatids and sperm to analyze the DiD+ cells and captured them for scRNA-Seq to test whether the cells we injected the week before differentiated into spermatids and sperm. The DiD+ cells were captured from whole-animal cell suspension produced from the reproductive recipient animals that reached again to stage B. *B. schlosseri* cells have natural autofluorescence in the far-red channel (used to detect DiD+ cells) and the DiD+ cells that were captured for scRNA-Seq from non-transplanted control animals were attributed to this autofluorescence.

For the second transplantation experiment, GSCs were sorted, labeled and transplanted as described above (n=3) while pigment cells were sorted according to a previous published protocol^22^, labeled and transplanted as a control (n=2). Seven days after transplantation, recipient testes were dissected and dissociated into single-cell suspensions. The proportion of DiD+ cells in each recipient was then analyzed by flow cytometry. Statistical significance was calculated using Welch’s t-test.

For the third transplantation experiment, GSCs and spermatozoa were sorted as described above. GSCs were labeled with DiD, whereas spermatozoa were labeled with DiO. Approximately 30,000 cells of each cell type were premixed and co-transplanted into the same recipients (n=4). Seven days after transplantation, recipient testes were dissected and examined by confocal microscopy (Zeiss LSM700) to identify transplanted cells. Cell islands, which exhibit high phagocytic activity and had engulfed large numbers of transplanted cells, were used as a positive control to confirm successful transplantation.

### Single cell RNA sampling and sequencing

To identify the GSCs, which represent a small fraction of the total cells in the mature testis and in the cell islands niche, we employed index sorting followed by the Smart-seq3 method^32^. This plate-based single cell approach provided paired-end reads and full-length transcripts, offering a higher sensitivity and coverage per cell that allowed us to identify a rare cell population (GSCs), which might not have been possible if we used droplet-based platforms for scRNA-Seq. Single cells were index-sorted using Sony MA900 FACS and processed using a modified Smart-seq3 protocol^32,72^. Cells were sorted into 96-well plates containing 2 μl of lysis buffer and plates were immediately centrifuged at a speed of 3000 G for 30 sec, snap frozen on dry ice, and stored at −80 °C. Reagents for all procedures including lysis plates preparation, reverse transcription, PCR amplification, plates clean-up and library preparation were used as previously described^72^. Before reverse transcription we thawed the cell-containing lysis plates on ice and incubated them at 72 °C for 3 min followed by immediate snap chill on ice to anneal the oligo dT_30_VN primer. Reverse transcription reagents (3 µl per well) were added using a Mantis liquid handler (Formulatrix), and plates were incubated in a C1000 Touch Thermal Cycler (BioRad) at 42° C for 90 min, and then 70° C for 15 min to terminate the reaction. For cDNA amplification, 7.5 ul of PCR mix was added to each well and cycled using the following program: 98 °C for 3 min, followed by 25 cycles of 98 °C for 20 sec, 67 °C for 15 sec, and 72 °C for 6 min, and then by a final elongation step of 72 °C for 5 min. Clean-up of amplified cDNA was performed using 0.65–0.75x volume of calibrated AMPure XP beads (Beckman Coulter) to remove residual reaction components and oligo fragments smaller than 400 base pairs, and eluted in 12.5 μl of ultra-pure water. The clean-up step was carried out using the Bravo automated liquid handling platform (Agilent). For quality control, 1 μl of purified cDNA was taken from each well. cDNA concentrations and size distributions were determined using Fragment Analyzer (Advanced Analytical). The wells within the 96-well plates were combined into new 384-well plates using the Mosquito X1 liquid handler (SPT Labtech). In the destination plate, the cDNA concentration was normalized, by diluting with ultra-pure water, to ensure it did not exceed 1 ng/μl.

Normalized cDNA was used to prepare Illumina sequencing libraries. Tagmentation was performed by combining 0.8 μl cDNA with 1.2 μl homebrew Tn5 mix as previously described^72^. The reaction was terminated by adding 0.4 μl neutralization buffer (0.1% SDS). Indexing PCR reactions were performed by adding 5 μM i5 and i7 index primers, 0.4 μl each (Integrated DNA Technologies, custom made 7680-plex unique dual index-primer set), and 1.2 μl KAPA HiFi HotStart ReadyMix (Kapa Biosystems). PCR amplification was performed using the following program: 72 °C for 3 min, 95 °C for 30 sec followed by 11 cycles of 98 °C for 10 sec, 67 °C for 30 sec, and 72 °C for 1 min. We pooled 1 μl of each well within the 384-well plates followed by a purification step using 0.8x volume of AMPure XP beads (Beckman Coulter). The 384-cell library pool from each plate was analyzed for concentration and size distribution using Fragment Analyzer (Advanced Analytical). All 384-cell library pools were normalized for concentration and pooled into a single tube that contained all the cell libraries. Following another purification and concentration step using 0.8x AMPure beads, the final library pool was sequenced on the NovaSeq 6000 S4 flow cell (Illumina) to obtain ∼1.5 million 2×150 base-pair paired-end reads per cell.

### Single cell RNA analysis

For read mapping, sequences were demultiplexed followed by quality trimming with TrimGalore v0.5.0 and aligned to our newly generated *B. schlosseri* genome with STAR program^73^ (parameters: --outFilterMultimapNmax 20 --outFilterMismatchNoverLmax 0.04 -- alignSJoverhangMin 8 --alignSJDBoverhangMin 1 --sjdbScore 1 --alignIntronMax 1,000,000 -- alignMatesGapMax 1,000,000). The bam files from the STAR alignment were used to calculate expression levels of genes and transcripts and incorporate them into gene count tables. Gene counting was executed using HTSeq v2.0.1_py39^74^. Gene count tables were combined with metadata using the Scanpy python package v.1.10.1^75^. We filtered out cells that expressed less than 5 genes. The data were normalized using size factor normalization so that every cell has 10,000 read counts, log transformed and scaled to a maximum value of 10. Highly-variable genes were computed using default parameters, and principle component analysis (PCA) was used to compute the neighborhood graph, followed by data clustering using the Leiden method^76^. Step-by-step instructions and detailed code to reproduce our data preprocessing and analysis are available in the supplementary information. To assist with cluster annotation, we used GeneAnalytics^77^ to identify the highest cell type score based on the top 200 most highly expressed genes in each cluster. For the *Botryllus* aging experiment, we assessed the potential contribution of technical variability between young and old samples by comparing standard scRNA-Seq quality-control metrics across groups, including the number of detected genes and total counts per cell. These analyses revealed broadly comparable distributions within both GSC and STD/SZ populations, indicating similar library complexity and sequencing depth between age groups before downstream comparative analyses (Extended Data Fig. 8). To increase the power of the analysis while accounting for potential plate-specific technical variation, we integrated, using Scanorama^78^, the datasets using the independently processed Smart-seq3 plate replicates as the batch variable.

### Human and mouse single cell RNA dataset collection and meta-analysis

Mouse single cell validation used 10X Genomics data from Zhang et al.^24^, with the data itself being downloaded from the China National Center for Bioinformation (NGDC) portal, with accession id OMIX1000 (https://ngdc.cncb.ac.cn/omix/release/OMIX1000). Data for each sample was downloaded individually and merged into a single object for analysis. Cells were filtered based on the same criteria as the original paper with cells with <300 total genes or >20% mitochondrial being excluded from the subsequent data analysis.

The human single cell validation used 10X Genomics data from Nie et al.^23^, with the data itself taken from the Gene Expression Omnibus (GEO) accession id GSE182786 (https://www.ncbi.nlm.nih.gov/geo/query/acc.cgi?acc=GSE182786). As with the mouse data filtering was done in the exact same manner as the original paper, in this case with cells with <800 expressed genes or >50% mitochondrial genes being excluded for further analysis. Extra sample information regarding smoking status, bmi and the exact age of each sample was taken from the paper’s supplementary Figure S1 A and Figure S6 G^23^.

For each dataset, cluster analysis and annotation were performed as described above for *Botyllus*. Following cluster annotation, the different cell types (GSC and STD/SZ) were grouped together and differential expression analysis for the different gene markers in each cell type was performed (Supplementary Tables 8 and 9). As for *Botryllus*, to compare expression of candidate gene markers in the mouse and human datasets, for each dataset, the merged data was loaded into SCANPY^75^ and figures were generated using the scanpy.pl.matrixplot function, with the gene expression of each of the genes being scaled (meaning that for each gene individually the sample with maximum expression would have a value of 1 and the sample minimum value for each gene being 0 and all other samples corresponding within that range). For each dataset, a matrix plot for the gene markers was generated based on the average of each age group and cell type (young and old GSC and young and old STD/SZ) and focused on the ability of the gene markers to separate the young and old samples similarly to how they could in the young and old *Botryllus* single-cell data generated as part of this study. For each dataset, an additional matrix plot separating the expression of the gene markers in the different cell types among individual biological samples was generated.

### Differential expression analysis

To further investigate differential expressions of the main genes discussed in this study (*DDX4*, *PIWIL1*, *SYCP3*, *H1F0*, *PSME3*, *UBE2R2*, *SAXO4*, *PRM1*, *SPA17*, *TSSK1B*) between young and old *B. schlosseri* cells involved in spermatogenesis, we used the gene count tables generated for the scRNA-Seq analysis. The data was normalized (Log2 CPM) and gene expression was compared across age groups and cell types for the selected genes and plotted using GraphPad Prism 10.3.1. Differential expression analysis was performed using Welch’s t-test followed by multiple correction analysis. Significance levels were set to 0.05*, 0.01** and 0.001***.

### Bulk RNA sampling, sequencing, and analysis

To validate the enrichment of GSCs from whole animals using our gating strategy, cells were sorted from the ALDH+ GSC gate, while control cells, for comparison, were sorted from the total live-cell population (Extended Data Fig. 5). Seven biological replicates derived from 4 different colonies were used to sort 100,000 cells from each gate (“GSC” and “Total”) for RNA sequencing. Cells were sorted directly into a 1.5 ml tube with 750 µL of Trizol-LS (Life Technologies) and RNA was extracted according to the manufacturer’s instructions. Briefly, the samples were chloroform-extracted, precipitated with isopropanol, washed twice with ethanol and dissolved in molecular-grade, RNase-free water. Samples were analyzed on an Agilent 4150 TapeStation to determine RNA integration and quantity prior to library preparation. cDNA and libraries were prepared using NEBNext^®^ Single Cell/Low Input RNA Library Prep Kit and barcoded using NEBNext Multiplex Oligos for Illumina. All magnetic bead purification was accomplished using DNA Spri Beads. Samples were then analyzed on an Agilent 4150 TapeStation to determine the concentration of each sample prior to sequencing. On average, 35 million 2×150 bp reads (NovaSeq X Plus) were sequenced for each library.

Gene count tables were generated as described above for the scRNA-Seq. Differential expression analysis was performed using Welch’s t-test followed by multiple correction analysis.

### Genomic DNA isolation

Genomic DNA (gDNA) was extracted from a whole system of a 3.5-year-old *B. schlosseri* colony (strain: 356a) that was collected in Monterey Bay, CA. This strain was also used for the assembly of our previously generated The *B. schlosseri* genome^25^ to simplify comparison between the gene models of the two different genome assemblies. The gDNA was prepared using a modified silica based DNA extraction technique^79^. The *B. schlosseri* sample was ground under liquid nitrogen to a fine powder, then transferred into a 1.5 ml tube containing a solution of 10 M Guanidine Isothiocyanate lysis buffer and GeneClean Glass Milk (MP Biomedicals). Lysed samples were incubated at 56 °C, washed and pelleted by centrifugation, then eluted in 0.1M TE. The final sample yielded 2.02 µg of high molecular weight gDNA with an average fragment size of 16 kb, which was quantified using Qubit (dsDNA broad range kit) and visualized using an Agilent Fragment Analyzer (genomic DNA 50 kb kit).

### Genome sequencing and assembly

A sequencing library was prepared according to PacBio’s “Preparing Whole Genome and Metagenome Libraries Using SMRTbell Prep Kit 3.0” protocol, excluding shearing and size selection to preserve gDNA input. The final library size was verified using an Agilent 4150 TapeStation and quantified with Qubit. The Sequel II Binding Kit 2.0 was used for primer annealing and polymerase binding of the library, followed by loading (140 pM) on two 8M SMRT cells and sequenced on PacBio’s Sequel IIe system. The sequencing “HiFi” application, circular consensus sequencing (CCS), and genome assembly were set up and performed in SMRT Link (v 11.0). The productivity rate (P1) for the two SMRT cells was 74%, on average, yielding a combined 4.9M HiFi reads with an average read length of 7.7 kb and Q38 read quality. SMRT Link’s genome assembly analysis (default settings) yielded 2.1K contigs with an N50 contig length of 1.1 Mb. This represents a substantial improvement over our original *B. schlosseri* genome assembly^25^, demonstrating the enhanced resolution achieved through long-read sequencing and assembly programs that were specifically designed for it.

### Genome annotation

We used raw sequence reads (NCBI accessions: PRJNA732987, PRJNA579844, PRJNA205369 and SRP022042) of previously published data^16,19,25,55^ to annotate the newly generated *B. schlosseri* genome. The raw sequence reads were analyzed using the NeatSeq-Flow platform^80^. The sequences were quality trimmed and filtered using Trim Galore (v 0.4.5) and cutadapt (v 1.15 parameters: --length 50 –quality 20 --max_n 1 --trim-n). Quality trimmed reads were mapped to our newly generated *B. schlosseri* genome. Mapping was done using STAR program^73^ (v 2.7.11a parameters: --outFilterMultimapNmax 20 –outFilterMismatchNoverLmax 0.04 --alignSJoverhangMin 8 --alignSJDBoverhangMin 1 --sjdbScore 1 --alignIntronMax 20,000 --alignMatesGapMax 20,000). The resulting alignment was used to create Genome guided transcriptome assembly using the Trinity program^81^ (V2.8.4 parameters: --min_kmer_cov 2 -- genome_guided_max_intron 20,000). The Genome guided transcriptome assembly was performed for each strain independently (944axByd196 and Sc109e), and then the created transcriptomes were merged into one fasta file to be used for downstream gene model prediction and annotation. The new *B. schlosseri* genome was masked using the funannotate mask program^82^ (v 1.8.15 parameter: -s “Tunicata” -m “repeatmasker”). The Masked genome was then used alongside the merged transcriptome and the raw reads to train the genes model with the funannotate train program (parameters: --max_intronlen 20,000 --aligners gmap). Next, we used funannotate predict program (using the following parameters: --busco_db metazoan -- max_intronlen 20,000 --repeats2evm --organism other) that takes the PASA^83^ gene models from the previous step to train Augustus^84^. Then we added UTR data to the predictions and fixed gene models that disagreed with the RNA-seq data using the funannotate update program (parameter: --max_intronlen 20,000). Finally, we added functional annotation to the genes model by running funannotate iprscan and funannotate annotate (parameter: --busco_db metazoan). In the funannotate annotate, we have changed the code of the step (annotate.py) to not restrict annotation by pident (percent identity) and to not exclude annotations that do not match the “CuratedNames” database (funnotate ncbi_cleaned_gene_products.txt file).

### Histology and 3D reconstructions

For histology, samples were fixed in 1.5% glutaraldehyde in 0.2 M sodium cacodylate and 1.6% NaCl buffer for 2 h. Samples were washed 3 times in 0.2 M sodium cacodylate and 1.6% NaCl buffer followed by post-fixation in 1% OsO4 in 0.2 M cacodylate buffer for 1.5 h at 4 °C. The samples were dehydrated, soaked in Epon and propylene solution at 37 °C, 45 °C, and 60 °C and then embedded in resin. Sectioning (1 μm thick) was carried out using a Leica ultramicrotome followed by staining with toluidine blue. For 3D reconstructions, chains of histological sections stained with toluidine blue were photographed using Leica DMR optical microscope. The Images were then aligned using Adobe Photoshop and the 3D anatomical model of each sample, based on image stacks, was created with Amira software (Thermo Fisher).

### In situ hybridization

Short in situ hybridization antisense DNA probes were ordered as lyophilized oligo pools from Integrated DNA Technologies and were resuspended in nuclease-free water to a concentration of 0.5 µM. For each gene, the probe was designed based on the split-probe design of HCR v.3.0^85^ using HCR 3.0 Probe Maker v0.3.2^86^ with adjacent B1 or B3 amplification sequences (Supplementary Table 10).

For in situ hybridization, *B. schlosseri* samples were incubated in fixation buffer (4% PFA in 1x PBS) overnight at 4□°C, followed by 100% methanol dehydration and storage at −20□°C for at least 24□h. The samples were gradually rehydrated in 75%, 50% and 25% methanol diluted with PBS (5 min for each step) followed by two washes in 1x PBS for 5 and 10 min, respectively. The samples were permeabilized for 30 min at room temperature in detergent solution (1% SDS, 0.5% Tween-20, 50□mM Tris-HCl (pH 7.5), 1□mM EDTA (pH□8), 150 mM NaCl). The samples were then washed 2 × 5 min with PTw (0.1% Tween-20 in 1x PBS) followed by post-fixation in 4% PFA for 25 min and washed again 3 × 5 min with PTw. The samples were prehybridized in hybridization buffer (Molecular Instruments) for 30 min at 37□°C. The probes were then added to the hybridization buffer at a final concentration of 0.02 µM and the samples were allowed to hybridize at 37□°C for overnight but no more than 16 h. Following hybridization, the samples were washed 2 × 30 min in probe wash buffer (Molecular instruments) at 37□°C and then for 5 min in 5x SSCT (5x sodium chloride sodium citrate (SSC), 0.1% Tween-20) at room temperature. They were then pre-amplified in amplification buffer (Molecular Instruments) for 30□min. Meanwhile, H1 and H2 components of the HCR hairpins B1 or B3 coupled to either 546 or 647 amplifiers (Molecular Instruments) were incubated separately at 95□°C for 90□s, cooled down to room temperature in the dark for 30 min and then pooled together before being added to the amplification buffer at a final concentration of 30□nM. The amplification was then performed at room temperature for overnight but no more than 16 h. The samples were subsequently washed 2 × 5 min and additional 2 × 30 min in 5x SSCT followed by incubation for 2 h in PBS containing 0.8:500 DAPI. Finally, the samples were washed 3 × 10 min in PBS, transferred to 50% glycerol for at least 1 h and then mounted on a glass slide for imaging.

Imaging was done using Zeiss LSM700 confocal microscope. For each sample, a series of optical sections were taken with a *z*-step interval of 3–5□µm. Multichannel acquisitions were obtained by sequential imaging and the confocal optical sections were processed using ImageJ v.1.54j^87^.

## Supporting information

Extended Data Figures and Instructions to use the Genome Browser

## Data availability

The sequencing data generated during this study is available in NCBI. The scRNA-Seq data is under accessions: PRJNA1250573, PRJNA1248000, PRJNA1247989 and PRJNA1247985. The bulk RNA-Seq data is under accession: PRJNA1516662. The new genome is under accession PRJNA1246227. Step-by-step instructions and detailed code to reproduce our data preprocessing and analysis are available in the supplementary information.

The new *B. schlosseri* genome browser can be accessed via the following link. A user manual explaining how to navigate and use the browser is provided in the supplementary information. https://share.nautilus.stanford.edu/new_genome/?session=local-2i7_lmeM1tYlc7-rFKDcO

## Code availability

The custom codes generated and used for analyses during this study are available at Zenodo: https://doi.org/10.5281/zenodo.23689623

## Acknowledgements

We thank Federico Caicci, Imaging Facility at Dept. Biol. University of Padova, for technical support in preparing 3D reconstructions; Noah Gordon for assembling the new *B. schlosseri* genome browser and server; Suyash Raj for helpful advices regarding the RNA-Seq analyses; Elle Koren Sedova for figure adjustments; Wan-Jin Lu and Stuart Thompson for the helpful and valuable discussions; Christopher Lowe for valuable discussions and for providing resources and support allowing the execution of this project; Linda Quinn and Tejaswitha Naik for laboratory management and support; the Chan Zuckerberg BioHub (CZ Biohub) for long and short-read sequencing processing and expertise; the Stanford Protein and Nucleic Acid (PAN) Facility for technical support.

## Author information

These authors contributed equally and share last authorship: Irving L. Weissman, Ayelet Voskoboynik

## Contributions

Conception and design: T.L., A.V. and I.L.W.; mariculture: K.J.I. and K.J.P.; flow cytometry and sorting: T.L.; confocal microscopy: T.L.; scRNA isolation and library preparation: T.L., C.A., T.G., D.D.L., R.S., B.F.O.G.; gDNA isolation: K.J.P.; gDNA and scRNA sequencing: A.M.D., M.M., N.F.N.; genome assembly: A.M.D.; transplantation assays: T.L. and K.J.I.; in situ hybridization: T.L., T.G. and K.J.I.; histology and 3D reconstructions: V.V. and L.M.; sequencing analysis and bioinformatics: T.L., Y.V., E.M., L.L., D.D.L. and D.S.; writing of manuscript: T.L., T.R., B.R., A.V. and I.L.W.; all of the authors reviewed, edited and approved the manuscript. A. V. and I.L.W. contributed equally to this work and share senior authorship.

## Corresponding authors

Correspondence to Tom Levy, Irving L. Weissman and Ayelet Voskoboynik

## Ethics declarations

The authors declare no competing interests.

