## Extended Data Figures and Instructions to use the Genome Browser for "Germline stem cell isolation, lineage tracing, and aging in a protochordate"

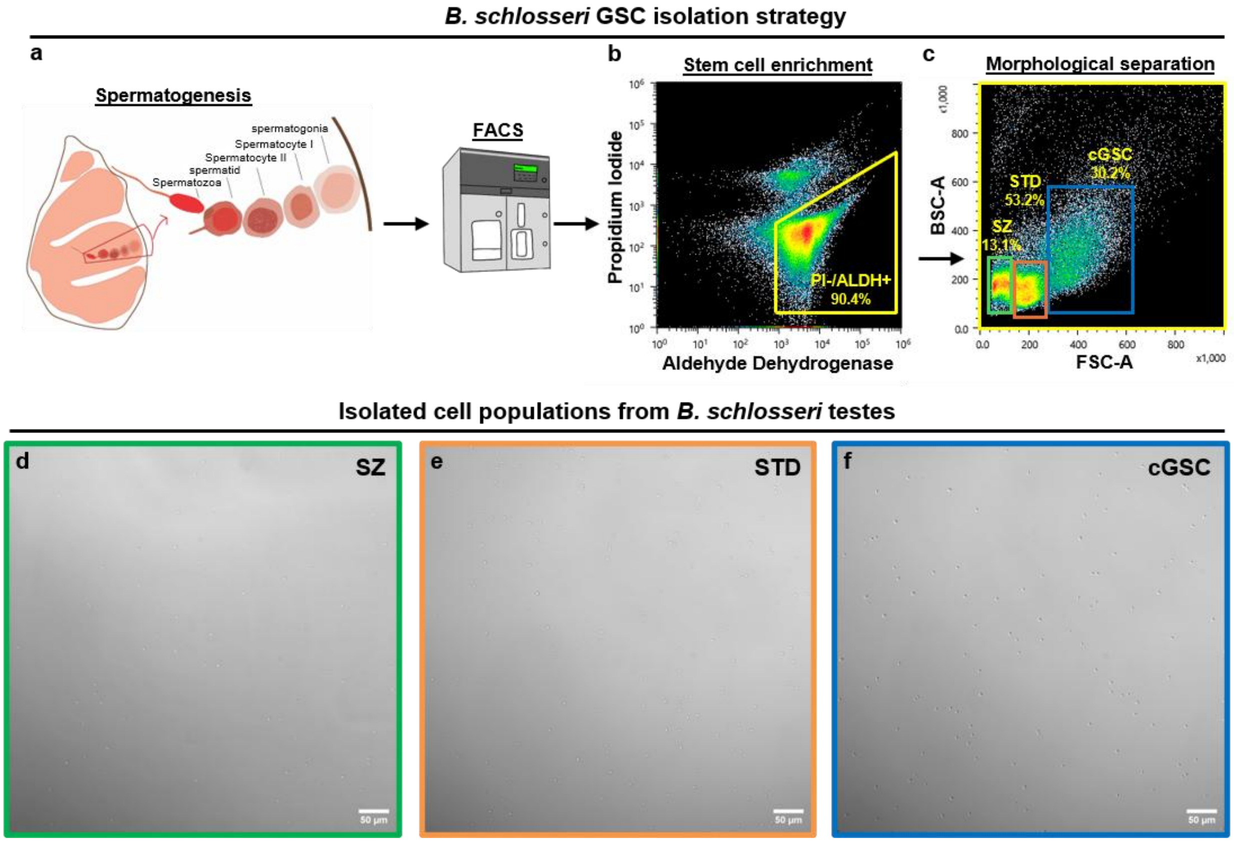


**Extended Data Fig. 1: ALDH stains the same cell populations as AP in the testis of *B. schlosseri*.**

**a,** Cell types identified during spermatogenesis. Spermatogonia are located at the lobe's periphery, followed by primary and secondary spermatocytes and spermatids (STD). Fully mature sperm (spermatozoa; SZ) are found in the lobe's center. **b,** Similar to AP+/PI- (Fig. 1), ALDH+/PI- cells from *B. schlosseri* testes were analyzed using FACS based on **c,** size (x-axis) and granularity (y-axis) and showed similar results. **d-f,** Representative brightfield images of cells sorted from the SZ **(d, green)**, STD **(e, orange)**, and candidate GSC **(cGSC; f, blue)** gates defined in **c**. Images show a wider field of view than Fig. 1 to illustrate the overall composition of the sorted populations.

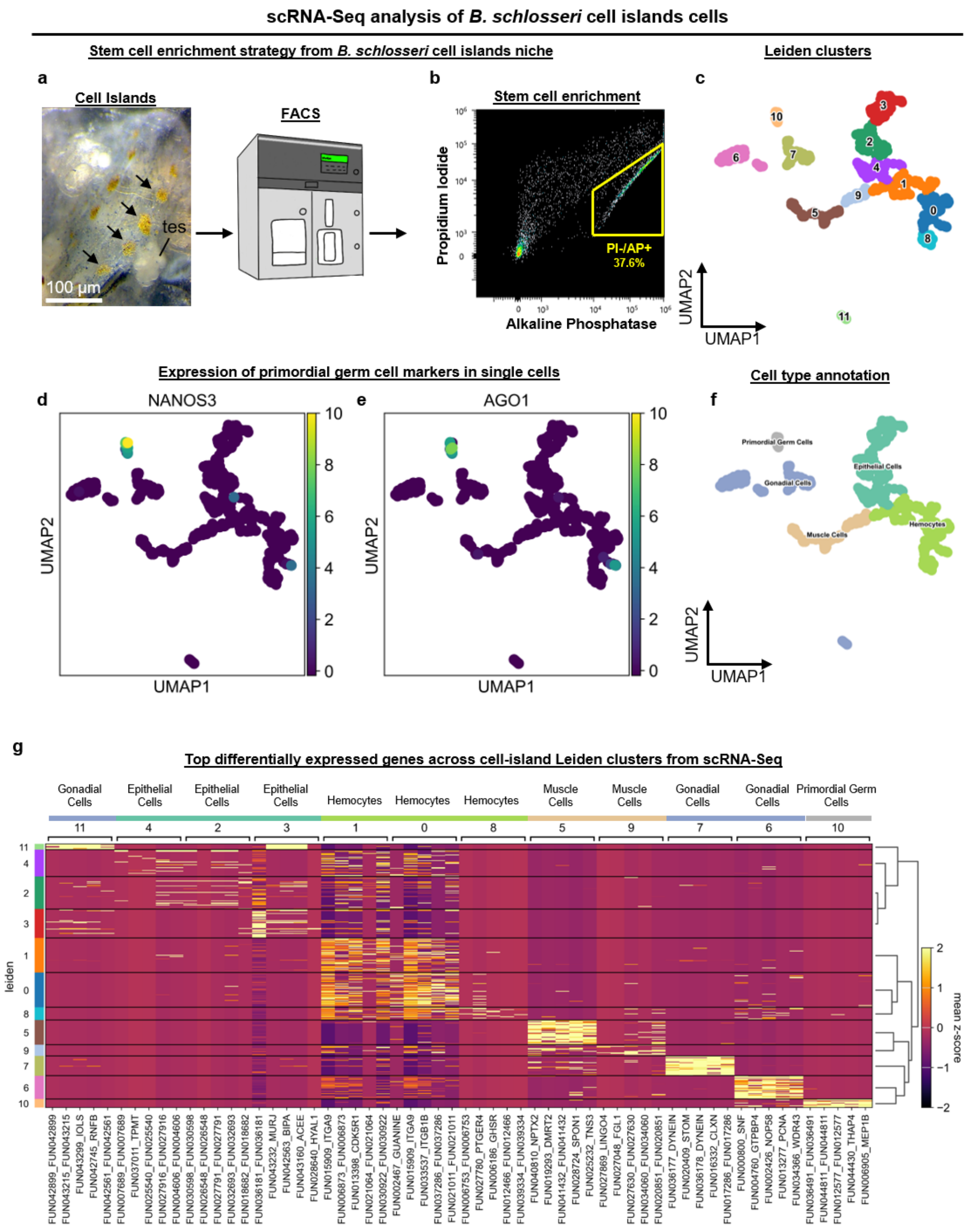


**Extended Data Fig. 2: scRNA-Seq reveals primordial germ cells in the cell islands niche.**

**a-c,** scRNA-Seq of index-sorted AP+/PI- cell-islands cells using Smart-seq3. **a,** Representative photo of an adult zooid. Cell-islands are labeled with black arrows and the testis (tes) is also visible. **b,** The AP+/PI- cell population that was sorted for sequencing. **c,** UMAP plot showing the Leiden clusters with cluster numbers. **d-f,** Cell type annotation. UMAP expression plots of primordial germ cell markers: *NANOS3* **(d)** and *AGO1* **(e). f,** UMAP plot with annotated cell type identities based on expressed transcripts of known genes **(g)**. Related to Supplementary Table 5. *Apparent PI signal in AP-high cells likely reflects AP-channel spillover.

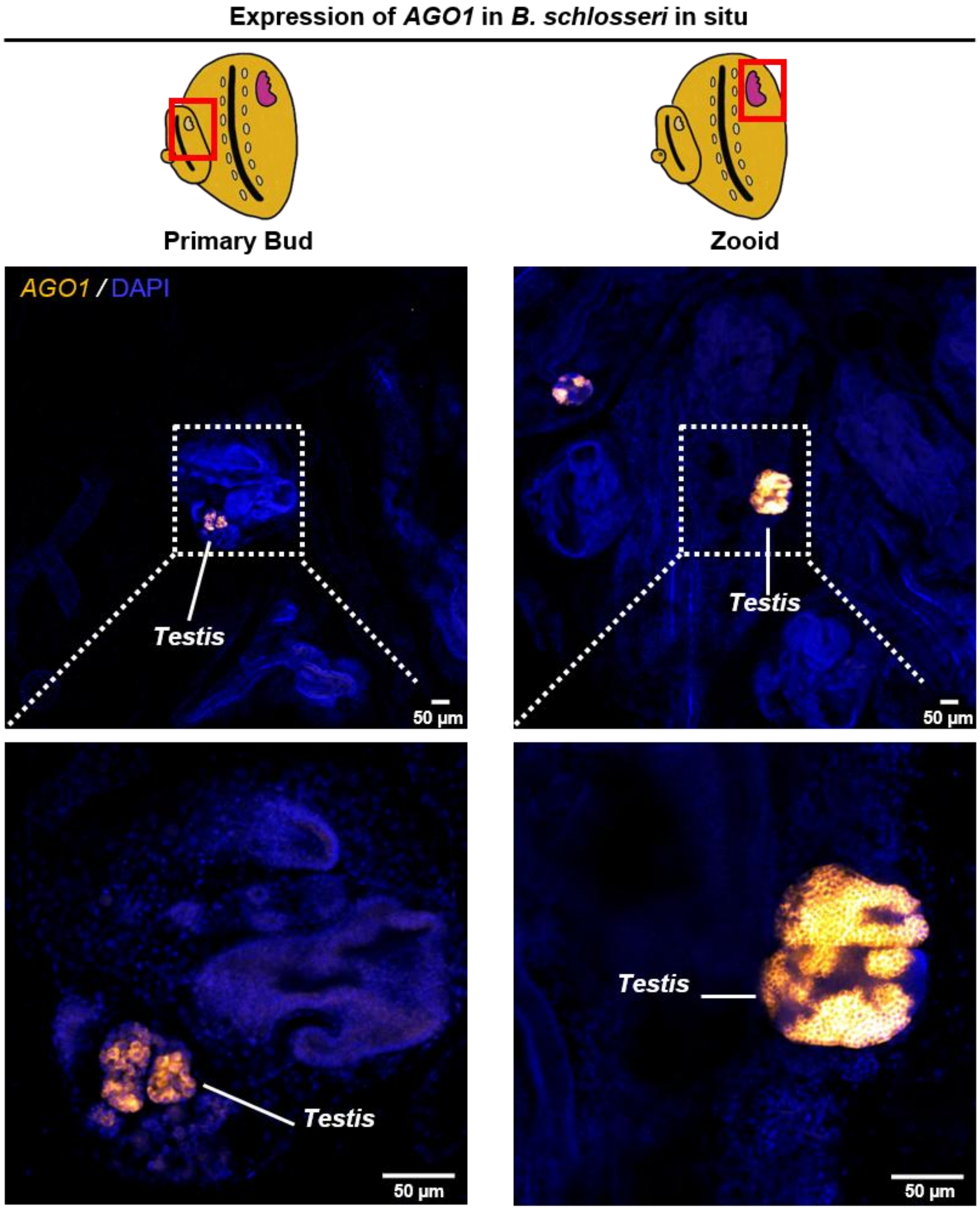


**Extended Data Fig. 3: *AGO1* is expressed in germline cells in *B. schlosseri* testes.**

HCR images showing *AGO1* (orange) expression in the developing gonad of both primary buds and adult zooids. Specimens are counterstained with DAPI (blue). Dashed boxes indicate the regions shown at the close-up images below. The testes in the primary bud and adult zooid are labeled with red frame in the schematic illustrations at the top.

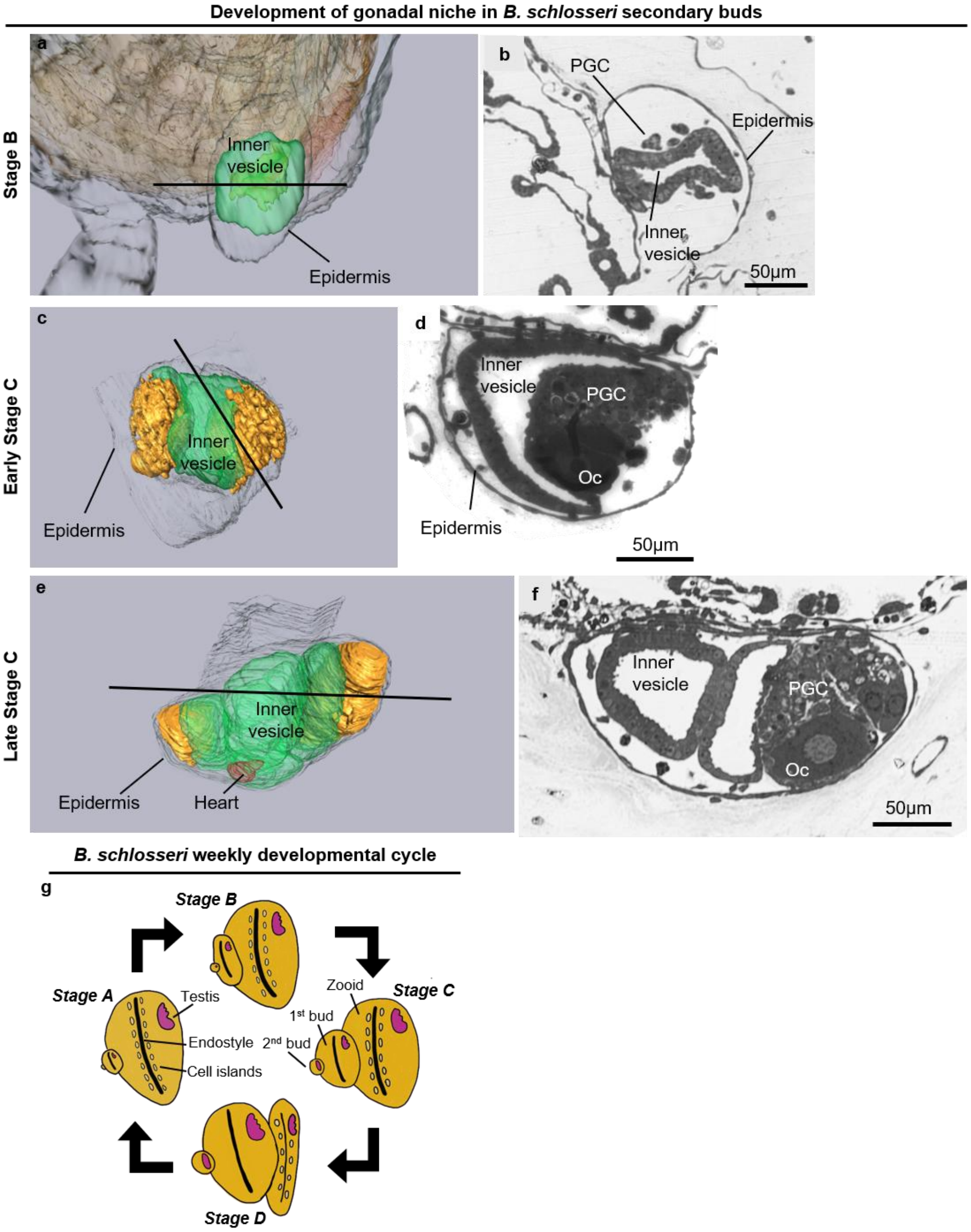


**Extended Data Fig. 4: Migration of primordial germ cells for gonadal development starts at the secondary bud.**

**a-f,** 3D reconstructions and cross sections of secondary buds with its inner vesicle (green) and developing gonad (orange) in a stage B colony **(a,b)**, an early-stage C colony **(c,d)** and a late-stage C colony **(e,f)**. The black line indicates the level of the section shown to the right of each 3D reconstruction. The bud's primordial germ cells (PGC), developing oocytes (Oc), inner vesicle, epidermis and heart are marked. **(g)** Schematic illustration of the *B. schlosseri* weekly developmental cycle.

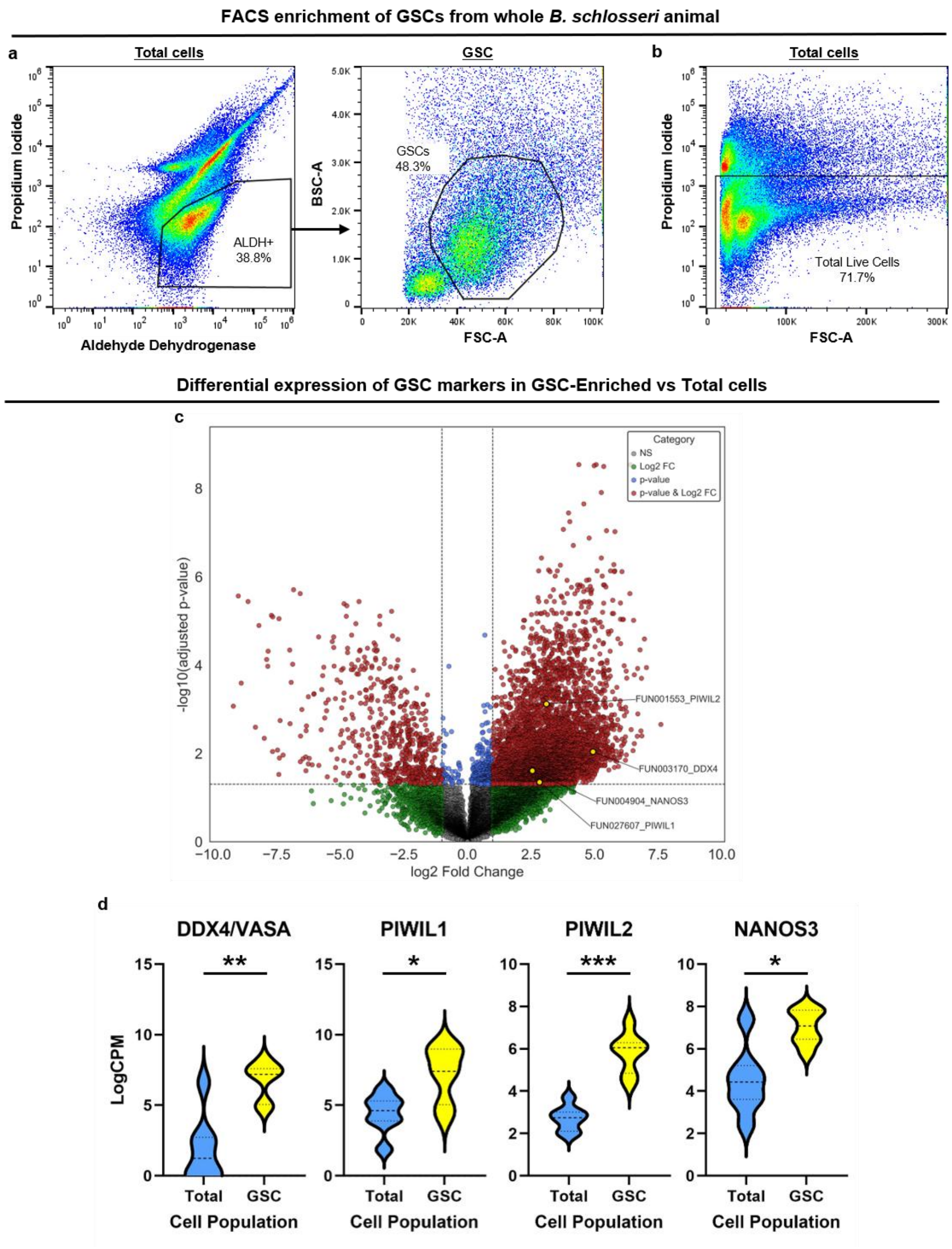


**Extended Data Fig. 5: FACS-based enrichment of GSCs from whole animals and transcriptomic validation by RNA-seq.**

**(a–b)** FACS gating strategy used to sort ALDH+ GSCs from whole-animal cell suspensions **(a)** and the total live-cell population **(b)** used for comparison. **(c)** Volcano plot of differential gene expression between sorted GSCs and total live cells, highlighting representative germline genes enriched in the GSCs in yellow (*DDX4*, *PIWIL1*, *PIWIL2*, *NANOS3*). Related to Supplementary Table 6. **(d)** LogCPM expression of the above representative germline markers in GSCs versus total live cells (RNA-seq results; n=7 for each group, *** $P<0.001$, ** $P<0.01$, * $P<0.05$; Statistical significance: two-tailed unpaired t test with Welch's correction is indicated; $P$ values were corrected for multiple testing using the Bonferroni method).

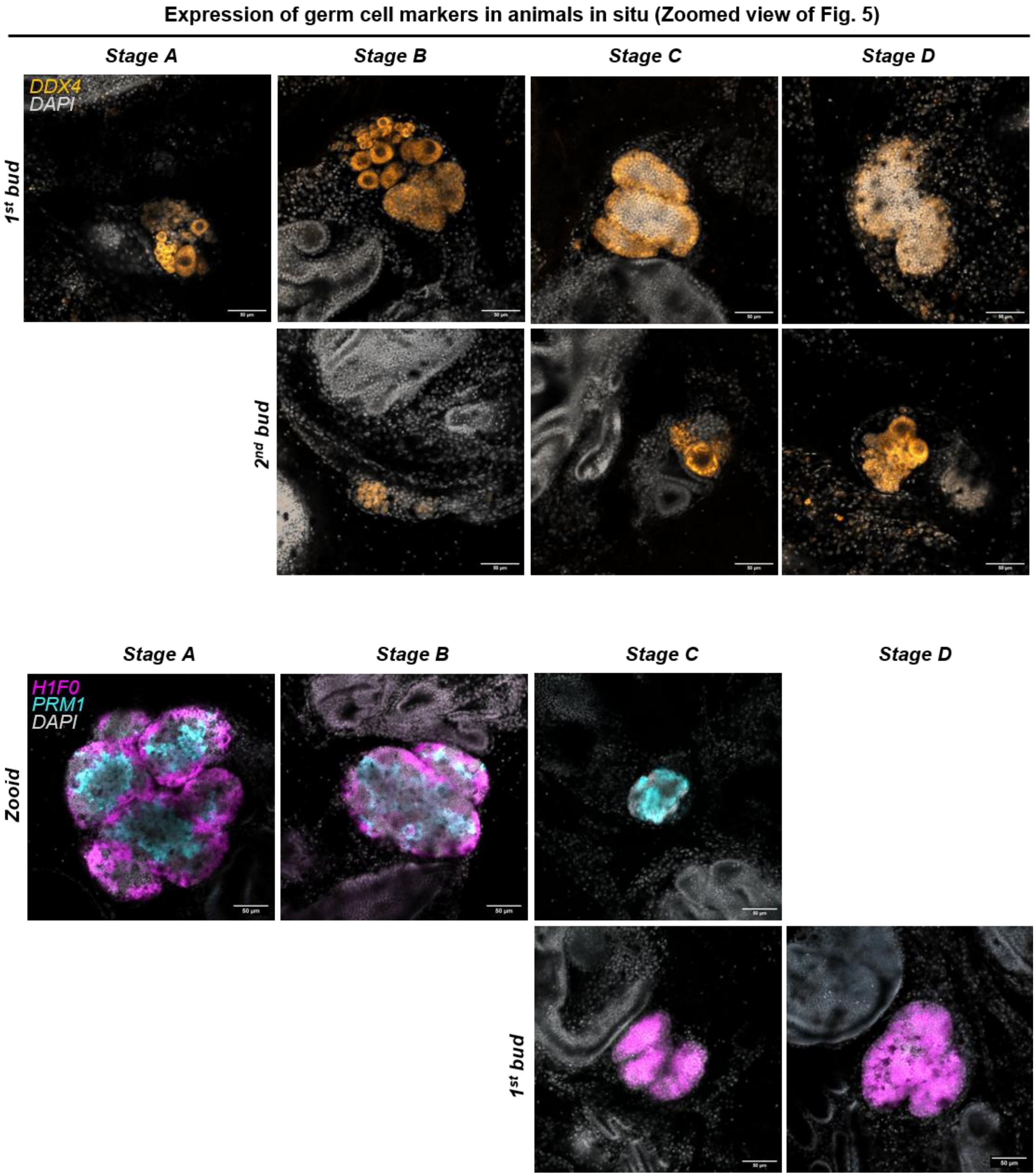


**Extended Data Fig. 6: Close-up views of HCR images shown in Fig. 5.**

Close-up images of the same specimens shown in Fig. 5, highlighting germ cell marker expression across developmental stages. Top: *DDX4/VASA* (orange) with DAPI (gray). Bottom: *H1F0* (magenta) and *PRM1* (cyan) with DAPI (gray).

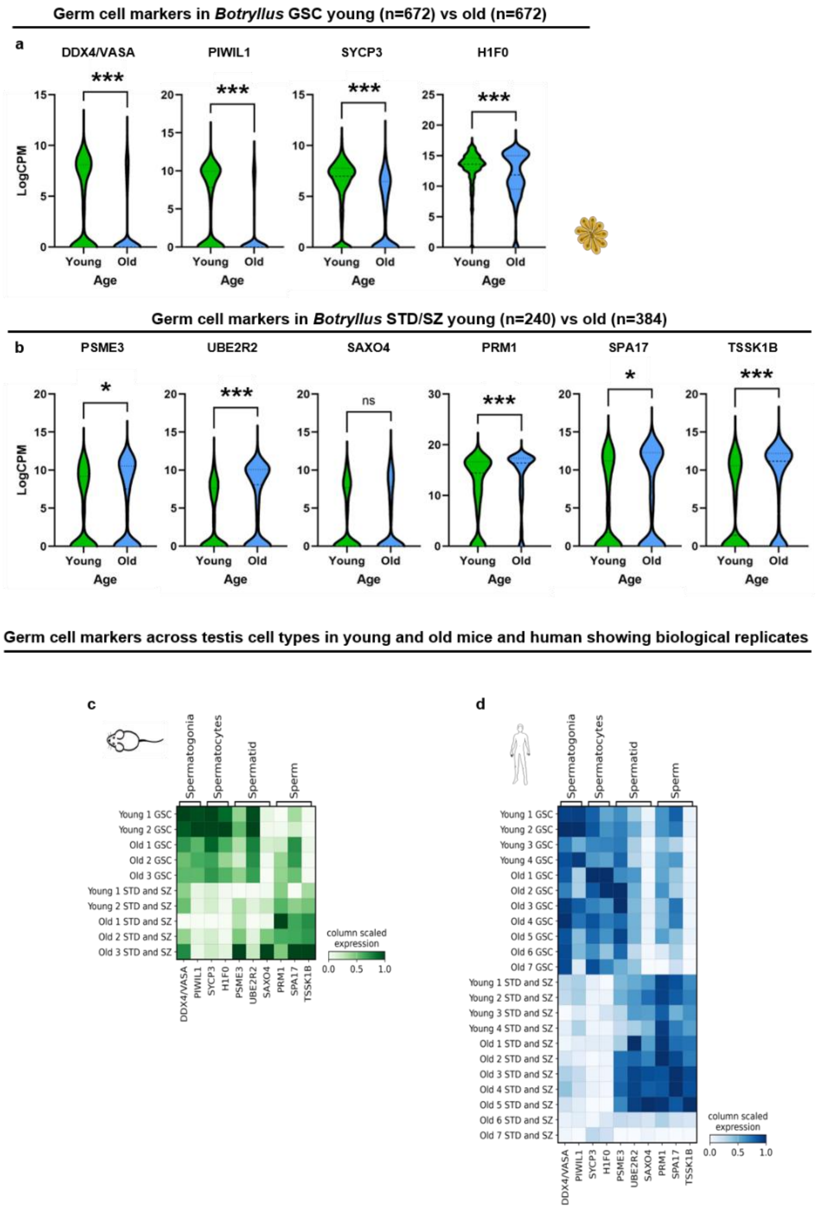


**Extended Data Fig. 7: Expanded analysis of age-associated germline marker expression in *B. schlosseri*, mouse and human.**

Violin plots showing scRNA-seq expression (logCPM) of representative germline markers used in this study (*DDX4, PIWIL1, SYCP3, H1F0, PSME3, UBE2R2, SAXO4, PRM1, SPA17, TSSK1B*) in young versus old *B. schlosseri* GSCs **(a)** and spermatids/sperm **(b)**. n indicates the number of single cells (Smart-seq3 libraries) analyzed per group. (*** $P$<0.001, ** $P$<0.01, * $P$<0.05; Statistical significance: two-tailed unpaired t test with Welch's correction is indicated; $P$ values were corrected for multiple testing using the Bonferroni method). **(c–d)** Expanded matrix plots underlying Fig. 6g,h, showing marker-gene expression across annotated germ cell types with biological replicates for *M. musculus* and *H. sapiens* (10x Genomics datasets from Zhang et al., 2023 and Nie et al., 2022).

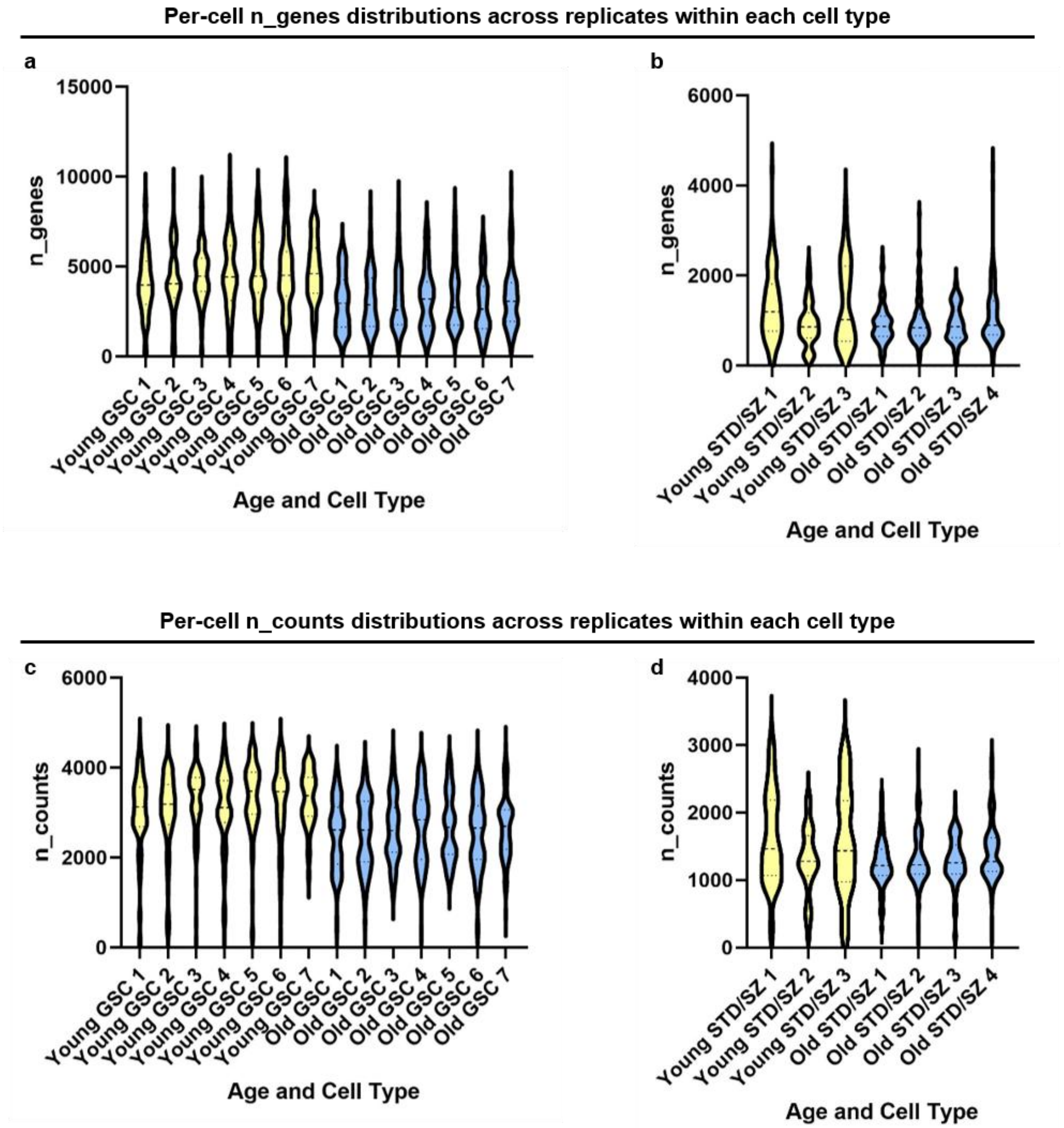


**Extended Data Fig. 8: scRNA-seq library complexity is consistent across colony-derived plate replicates within each age group.**

Violin plots show the distribution of the number of detected genes (n_genes) and total counts (n_counts) per cell across independently processed 96-well Smart-seq3 plate replicates from young and old *B. schlosseri* GSCs **(a,c)** and spermatids/sperm **(b,d)**. For each age group, cells were pooled from a colony comprising 4-6 systems and 6-10 zooids (biological units) per system prior to FACS sorting, providing broad biological sampling per replicate.

### <u>Manual for using the *Botryllus* Genome Browser (by Levy et al.)</u>

If you get the following error message – Refresh the website

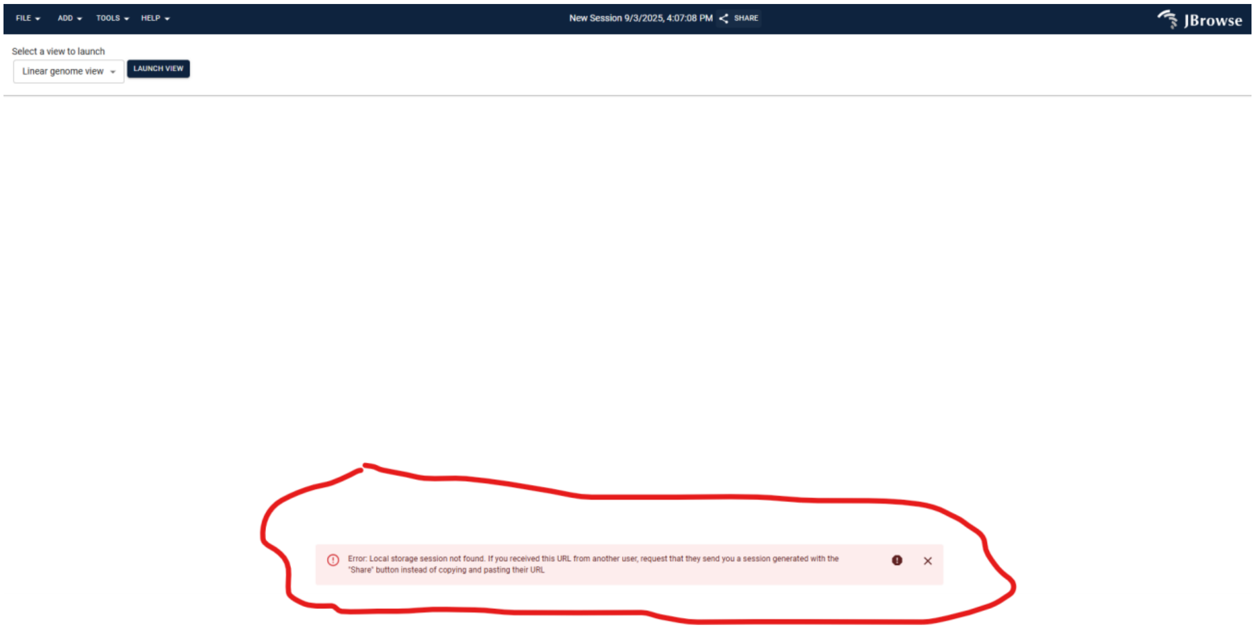


Click "Launch Review" > "Open" > "Open Track Selector"

This is the screen you will see:

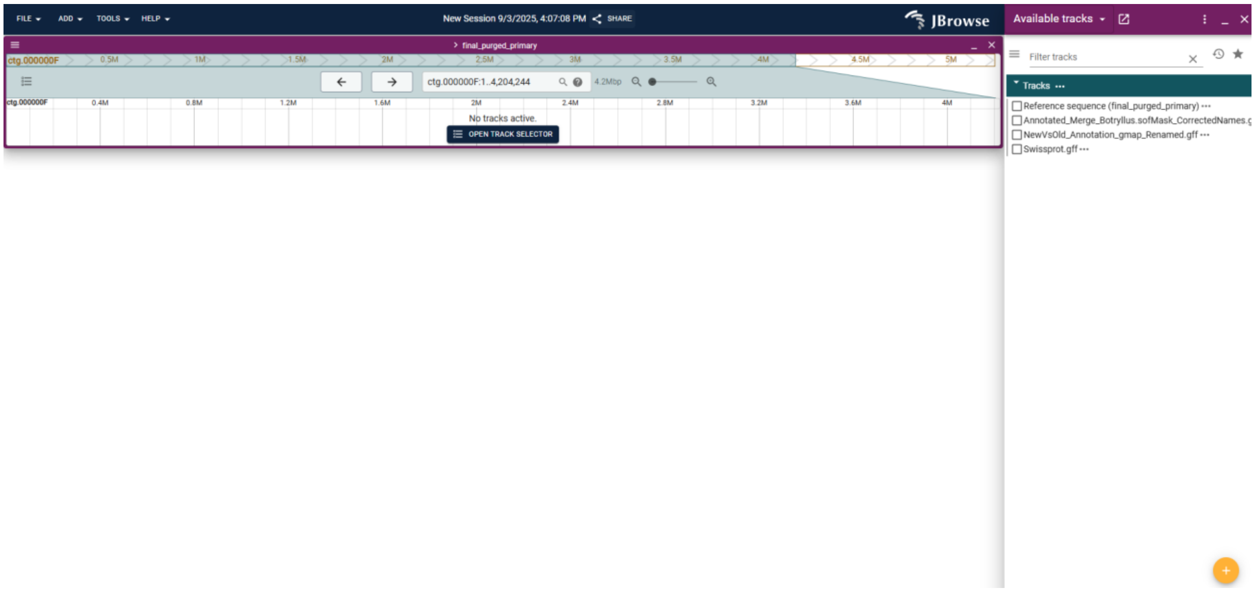


On the right side of the screen, you can select the following tracks:

"Reference Sequence" – The full *Botryllus* genomic sequence

"Annotated_Merge_Botryllus…" – The gene annotation we generated for the genome

"NewVSOld…" – The gene annotation based on Voskoboynik et al., *eLife*, 2013 (available for comparison)

"Swiss Prot" – Gene annotation based on Swiss Prot

To access a specific gene, simply type its name in the search box:

For example: "DDX46":

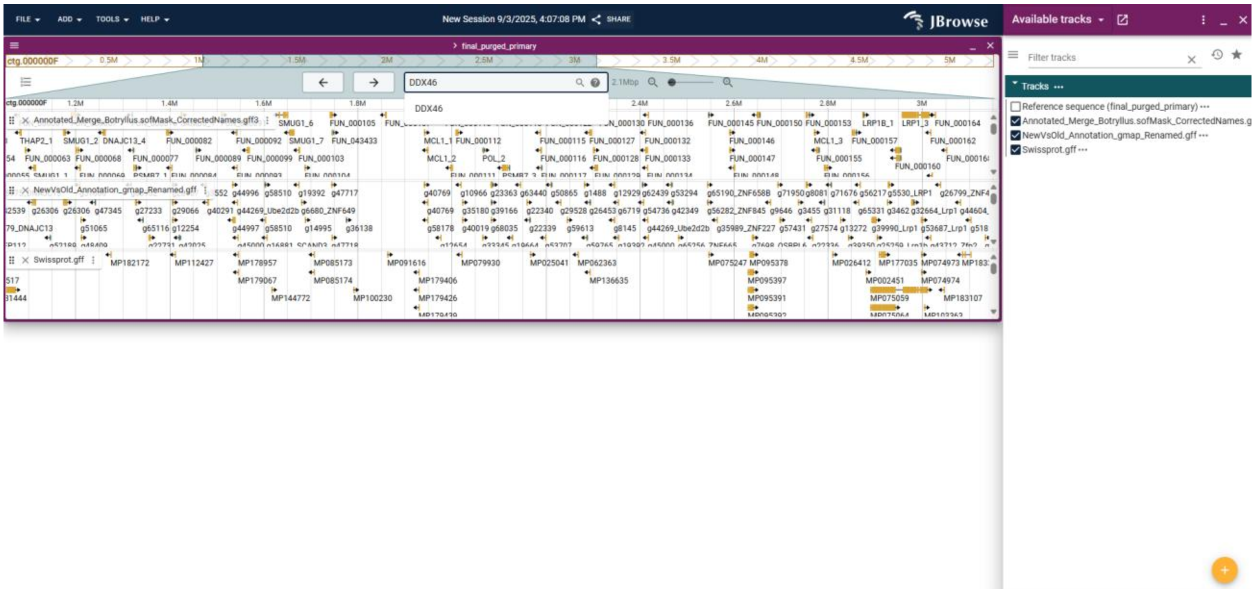


Select to see it in the new annotation by pressing "GO":

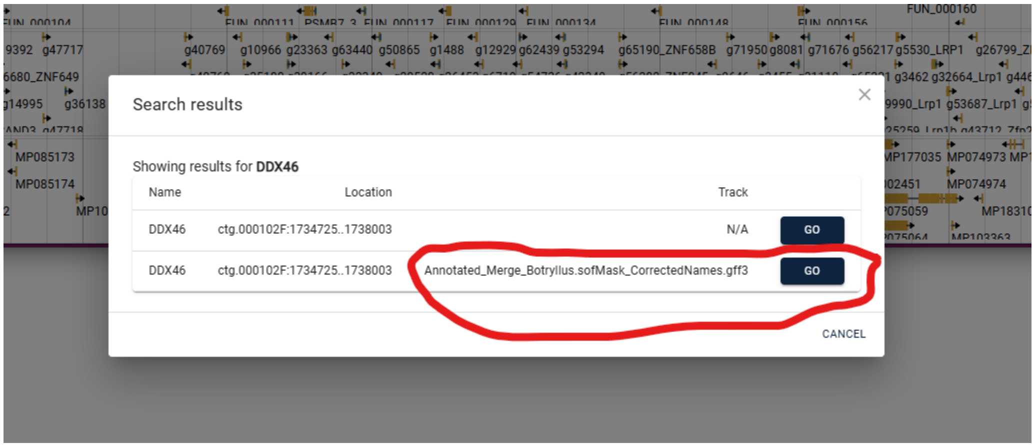

Click the yellow region of the gene to display all associated details, including its sequence, on the right side of the screen:

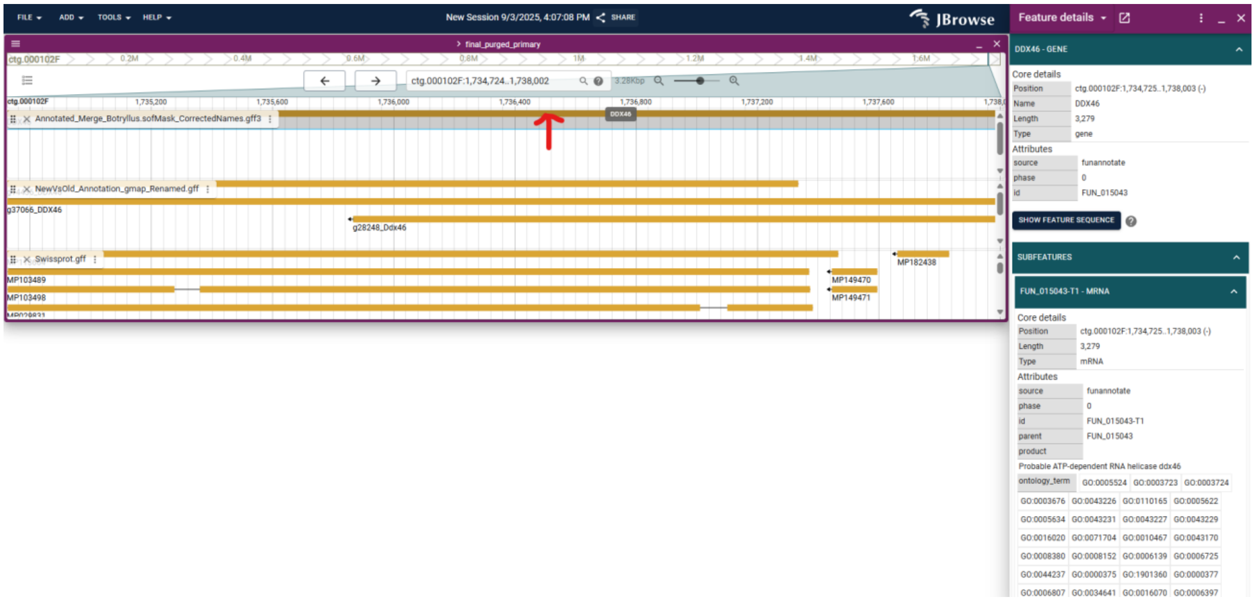
